# Associative memory retrieval modulates upcoming perceptual decisions

**DOI:** 10.1101/186817

**Authors:** Aaron M. Bornstein, Mariam Aly, Samuel F. Feng, Nicholas B. Turk-Browne, Kenneth A. Norman, Jonathan D. Cohen

**Affiliations:** Department of Cognitive Sciences, The University of California, Irvine, Irvine, CA, USA; Center for the Neurobiology of Learning and Memory, The University of California, Irvine, Irvine, CA, USA; Department of Psychology, Columbia University, New York, NY, USA; Department of Science and Engineering, Sorbonne University Abu Dhabi, Abu Dhabi, UAE; Department of Psychology, Yale University, New Haven, CT, USA; Department of Psychology and Neuroscience Institute, Princeton University, Princeton, NJ, USA

## Abstract

Expectations can inform fast, accurate decisions. But what informs expectations? Here we test the hypothesis that expectations are set by dynamic inference from memory. Participants performed a cue-guided perceptual decision task with independently-varying memory and sensory evidence. Cues established expectations by reminding participants of past stimulus-stimulus pairings, which predicted the likely target in a subsequent noisy image stream. Participant’s responses used both memory and sensory information, in accordance to their relative reliability. Formal model comparison showed that the sensory inference was best explained when its parameters were set dynamically at each trial by evidence sampled from memory. Supporting this model, neural pattern analysis revealed that responses to the probe were modulated by the specific content and fidelity of memory reinstatement that occurred before the probe appeared. Together, these results suggest that perceptual decisions arise from the continuous sampling of memory and sensory evidence.

---

Laboratory studies of decision-making tend to focus on choices made on the basis of a single kind of information – such as anticipated utility [1], sensory input [2], or mnemonic evidence [3, 4] – taken alone. But in the real world, our decisions depend on integrating information available from many sources – both external, such as visual, and internal, such as our memories.

For instance, when traveling on an unfamiliar train route, I might miss my intended stop. How do I figure out where to make the transfer to get back on my desired route? I could rely solely on sight – as the train stops at each station, quickly scan the platform for helpful signs or markings. I could rely solely on my memories – which station is next? Will it have the transfer I need? Both kinds of information can be unreliable: station platforms may look very similar, with distant or unhelpful signage, and my memories of the sequence of stations could be sparse or inconsistent (for instance, due to past detours). More likely, I will combine both kinds of information: query my memories about which stations might have transfers, and combine those with what I can see from a quick look out the door at each stop. By combining what I remember with what I see, I can improve my ability to figure out where I am – and, thus, what actions I should take.

A similar and open question in the laboratory study of perceptual decisions is how expectations should, and do, influence the inference process. Within the canonical sequential sampling framework [2, 3], expectations can be encoded as a change to either the starting point of the inference process, or the rate at which evidence is sampled [5–10]. Another, related idea is that expectations can dynamically impact both the rate and direction of sampling, with increasing influence as a decision takes longer to resolve [11]. However, all of these approaches assume that the content of expectations is fixed before the decision starts, whether by learning or by instruction. In the train analogy, the map is known with certainty, though the reliability of the visual cues vary from station to station (trial to trial).

Extensive work supports the idea that people query associative memory as an informative, though potentially unreliable, source of information for action selection [12]. Specifically, it has been shown that choice patterns and response time distributions during timed memory retrieval tasks are well-matched by the predictions of the drift-diffusion model (DDM; Ratcliff [3]), and that decisions can be made on the basis of sampled memories, similar to the way in which samples of visual input are used to guide perceptual decisions [13–17]. Further, hippocampal activity, as measured using fMRI, during decision-making appears to scale with the uncertainty of next-step associations [13, 18]. This finding concords with a broad literature supporting the involvement of the MTL, long associated with memory encoding and retrieval, in perceptual decisions [19]. Building on these results, we test the hypothesis that, during a cue-guided perceptual decision task, both types of evidence sampling occur as a single, continuous, inference process, with actions selected on the basis of the combined evidence.

Our hypothesis yields two main predictions. First, evidence sampling in such a task should begin before the onset of sensory information, with dynamics that change when the probe is presented. Specifically, before the probe, sampling should reflect the contents of memory retrieval and its consistency; after, the rate of sampling should be influenced by the coherence and content of visual information. Second, the hypothesis predicts that the sampling process is integrative across modalities. Specifically, decisions made after the onset of the probe should reflect the content and consistency of memories retrieved before the onset of the probe – that is, to the degree that the retrieved memories concord with visual samples, then the decision should be faster (and, conversely, when they disagree, responses should be slowed).

To test these predictions, we developed a memory-guided perceptual inference task. In the task, two distinct kinds of information – memory and sensory – indicated the correct response for that trial, and were made available at separate times. First, participants learned, by experience, a small set of cue-photograph pairs. Fractal cues were followed in quick succession by one of two face or house photographs, one more often than the other. Then, in the main phase of the task, these cues were used to establish expectations for a sensory decision. Specifically, a fractal cue triggered memories of the (face or scene) photographs that had been previously observed to follow in time. These memories served as evidence about the likely identity and reliability of a subsequent noisy visual probe stimulus – a rapidly alternating stream of photographs, one of which was the one predicted by the cue. Critically, participants could choose to respond at any time, including before the probe appeared. Therefore, their responses could reflect the influence of memory or sensory information alone, or some combination of the two.

We formalized our predictions using a multi-stage sequential sampling model that allows for dynamic changes in the rate – and, by implication, the content – of sampling [20–22]. In our task, the first stage of the model samples evidence from memories triggered by the fractal cue. The second stage carries forward the evidence sampled during stage one, while incorporating new samples, this time visual input from the noisy probe. This approach differs from previous models of expectation-guided perceptual inference in that it constructs expectations dynamically for each trial, using the cue to effectively anticipate the content of the probe. As a result, what the model “expects” will vary between decisions, depending on what evidence was sampled during the first stage.

Experiment 1 is a behavioral study that tests the first prediction of the model: that choices and response times in the task reflect a continuous inference process in which the rate of sampling changes with the onset of visual information. We fit our model to these data, and contrast its fit with that of non-continuous alternative models. Experiment 2 is an fMRI study that tests the second prediction: that evidence sampling from memory is itself a dynamical process, that evolves over the period prior to presentation of the cue, and the result of which is carried forward and affects the sensory inference process. We used Multivariate Pattern Analysis (MVPA) to measure, on a trial-by-trial basis, neural signatures of the degree and content of memory samples, and test their relationship to responses made after the onset of the flickering probe.

Taken together, the results of these experiments provide a new account of perceptual decisions, by demonstrating a critical role for integrated, dynamic inference from mnemonic, as well as sensory, information.

## Results

Participants performed a cue-guided perceptual inference task (Figure 1), in which fractal cues could be used to anticipate the content and coherence of a noisy probe stimulus that appeared after a short, variable-length delay (Figure 1b) The task encouraged participants to rely on evidence from memories, triggered by fractal cues, that could be consulted during the anticipation delay, and, after the delay, from a noisy visual probe: a stream of rapidly alternating photographs “flickering” at one of two levels of coherence (Figure 1c). The participant’s task was to press the key corresponding to the photograph that was most often present in the flickering probe. This photograph was referred to as the “target.” Critically, because the task was blocked such that each stimulus category corresponded to one coherence level in each block (Figure 1b), the fractal cue provided participants with two pieces of information about the probe: 1. the likelihood of each photograph being the target; and 2. the coherence of the flickering stream.

**Figure 1.**
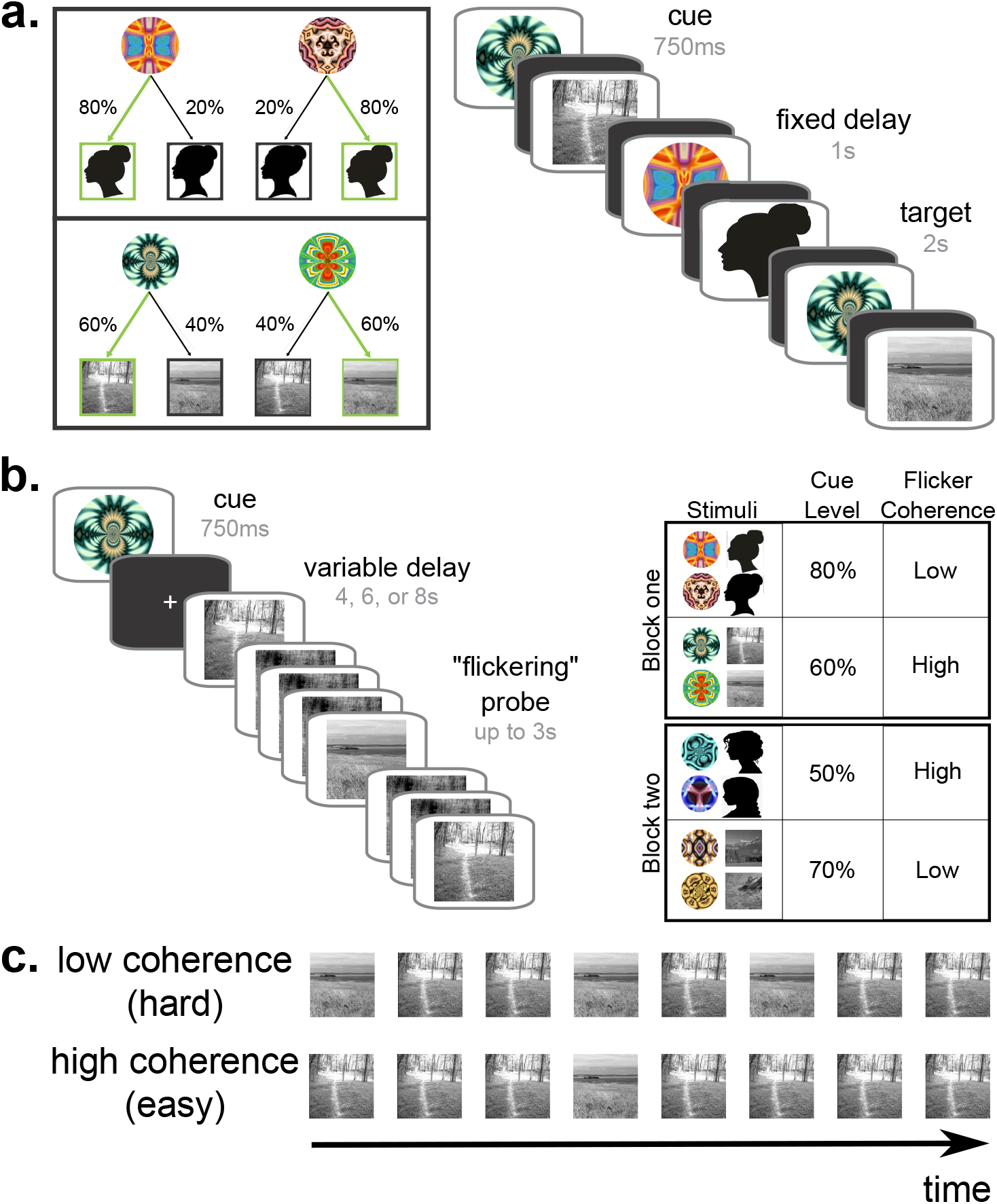
Cue-guided perceptual inference task. **(a)** In the *Sequence learning* phase, participants saw a series of 100 cue-target pairings. Each fractal cue was shown 25 times, randomly interleaved, and at each presentation followed by a target picture of either a face or a scene. They were told only to press the key associated with the target photograph, once it appeared. Cues were followed by photographs from one category only, and to each of the two pictures within that category according to complementary proportions (50%/50%, 60%/40%, 70%/30%, 80%/20%). This set of experiences of cue-target pairings provided associative memory traces that served as evidence samples for the task in *Test* phase. **(b)** In the *Test* phase, participants were again shown a fractal cue, but in this case the cue was followed by a “flickering” series of rapidly alternating pictures. Each frame of the series contained one of the two pictures from the cued category, or a phase-scrambled superposition of the two pictures. One picture, the *target*, was shown in the stream more often than the other. Participants were asked to respond by pressing the key associated with the target picture – critically, they could respond at any time after the onset of the fractal cue. In each block, each category was associated with a particular level of cue predictiveness, and flickering stimuli were either of high or low coherence – therefore, the fractal cue signaled the reliability of both memory and sensory evidence on that trial. **(c)** The flickering probe stimulus was calibrated to one of two levels of difficulty. In an earlier phase of the task, a staircasing procedure was used to determine the proportion of frames that would elicit either higher (85%) or lower (65%) accuracy on a neutral-prior (50/50) version of the flickering task. In this way, the flickering stream parametrically varied the weight of visual evidence favoring the target.

### Experiment 1

We first tested whether choices and response times reflected the influence of both memory evidence – operationalized via cue probability – and sensory evidence — operationalized via the coherence of the flickering probe. According to our hypothesized two-stage inference mechanism (Figure 2a), participants would respond more quickly and accurately when: 1. the cued memories more reliably predicted the identity of the upcoming photograph; 2. the observed visual evidence was more coherent; and 3. the cue predictions matched the visual evidence.

**Figure 2.**
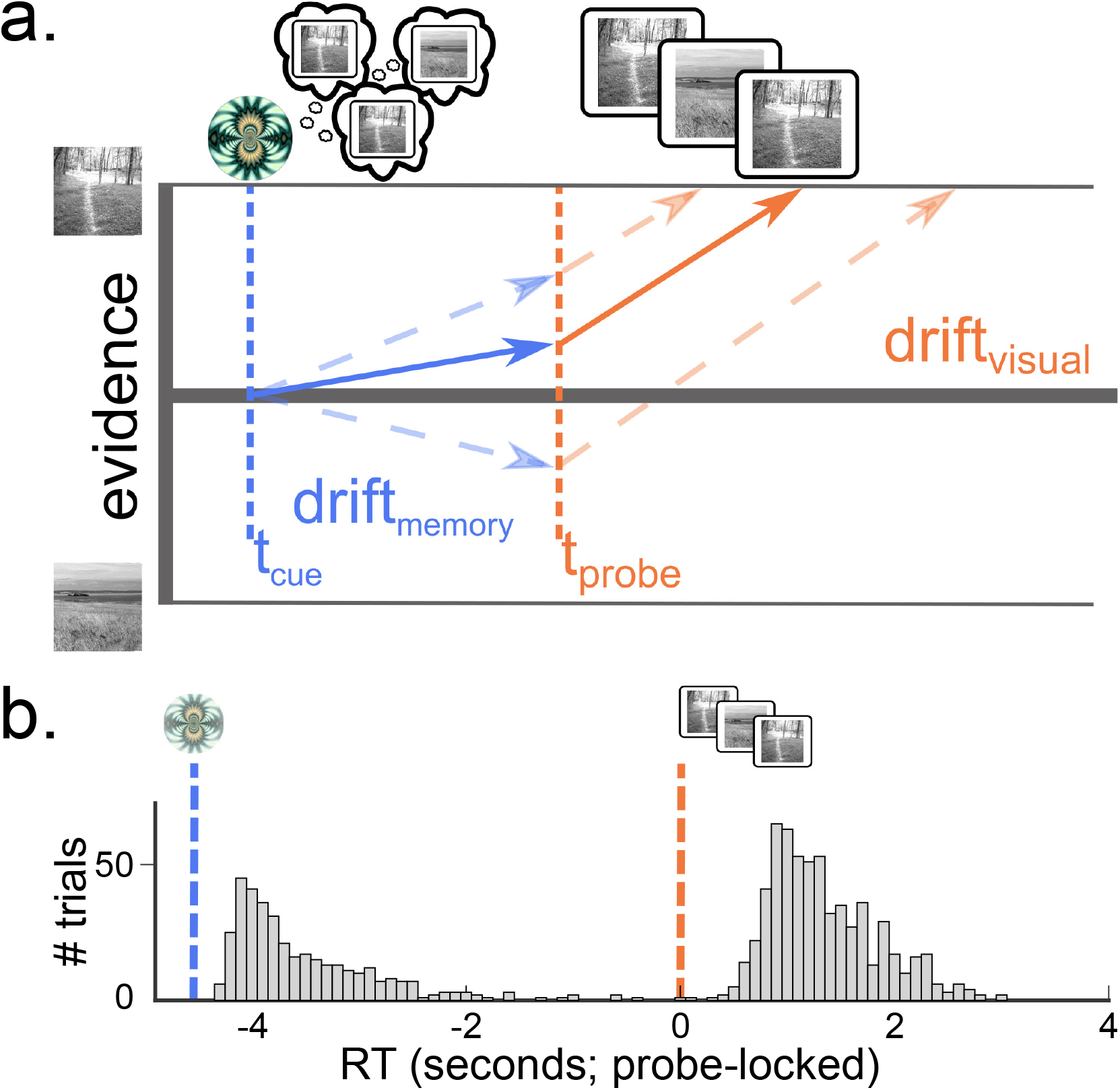
(a) Two-stage sequential sampling model. The *Multi-Stage Drift-Diffusion Model* (MSDDM) describes an sequential sampling process with time-varying drift rate [21]. The first drift rate corresponds to the period following onset of the fractal cue and preceding the onset of the flickering stream, while the second corresponds to the period after the onset of the stream. Critically, at each trial, the starting point of the second stage depends on the trajectory of the evidence sampling that occurs in the first, on that trial. **(b) Participants responded to both the fractal cue and the flickering probe**. Shown is a histogram of (probe-locked) response times on test-phase trials during the four-second delay condition. RT counts are aggregated across trials and participants, and binned in increments of 100ms. Separate peaks follow the onset of the fractal cue and the onset of the flickering stream, reflecting the fact that participants made responses on the basis of both types of information.

### Response times and accuracy

#### Responses reflect the influence of memory and sensory evidence

Consistent with a two-stage integration process, response times were distinctly bimodal, with separate peaks following the onsets of the fractal cue and the flickering stream (Figure 2b; RT distributions multi-modal within each ISI condition by Hartigan’s Dip Test [23]: all HDS*≥* 0.028, all *P* < .001).

Overall, participants responded accurately, matching the target photograph on 75.20% (SEM 0.085%) of trials (including only trials for which there was a “correct” response possible before stimulus onset – i.e. for cue levels 60%, 70%, 80%). This proportion was reliably greater than chance for all blocks individually (all *P ≤* 0.047 by binomial test of the proportion of correct responses within each block against the 50% chance level). Accuracy increased with both cue predictiveness (*R* = .195, *P* = .009 by bootstrap across participants; Figure 3a) and target coherence (*t*(27) = −4.430, *P* < .001 by two-tailed, paired two-sample t-test tested for the 28 participants who performed at least one block in which there was a predictive cue for both coherence conditions). Thus, as expected, both factors appeared to influence the decision making process.

**Figure 3.**
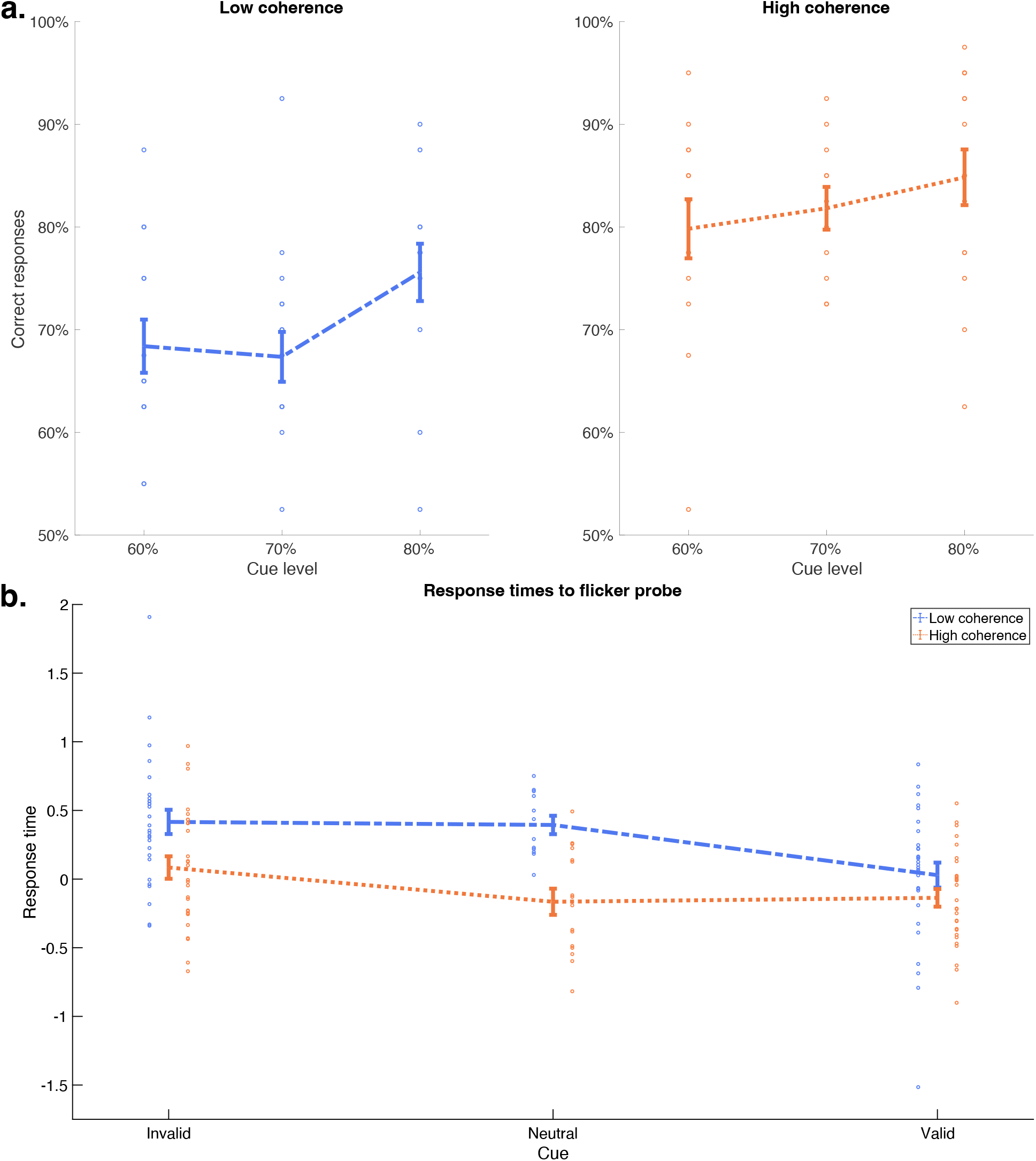
Choices and response times are modulated by the content and quality of both memory and sensory evidence. **(a) Memory and sensory evidence affected choice accuracy.** Across all trials, accuracy increased with both the frequency of the cue-target pairing (*R* = .195, *P* = .005) and the coherence of the flickering stream (*t*(27) = −4.430, *P* < .001). Cue-target association was a slightly stronger predictor of accuracy when sensory evidence was of low coherence (low coherence: *R* = .226, *P* = .019; high coherence: *R* = .180, *P* = .128; difference: *d* = 0.394). **(b) Response times to the flickering probe reflected a mixture of memory and sensory evidence**. When participants responded after the onset of the flickering stream, (probe-locked, z-scored) response times were lower when the target was the one that had been more often seen following the cue in the Learning phase (*R* = − .017. *P* = .044). The influence of the cue-target association was much stronger when the flickering stimulus was of low coherence (low coherence: *R* = −.147, P < .001; high coherence: *R* = −.065, *P* = .033; difference: *d* = 1.675). (Error bars are ± 1 SEM, across participants.)

Participants appeared to use the fractal cue to decide whether or not to respond “early” (before the onset of the probe stimulus). If the decision to respond early was driven by sampling from cued associative memory reinstatements, then it should be modulated by: 1. the quality of memory samples relative to sensory samples; and 2. the time available to sample. In other words, participants should have relied more on the cues when the associated memories were more consistent, when the cues signaled that the upcoming sensory evidence would be of low coherence, and when the anticipation delay was longer.

Consistent with this model, the proportion of early responses increased with the predictiveness of the fractal cue (*R* = .222, *P* < .001), and this relationship was driven by trials on which the cue signaled that the perceptual stimulus would be of low coherence (for low coherence trials, the correlation between cue predictiveness and early responses was *R* = .366, *P* < .001; for high-coherence trials it was *R* = .087, *P* = .161; difference: *d* = 3.907). Further, these responses were faster when memory evidence was stronger. Within the group of early responses, RTs showed a trend towards being faster as the fractal cue – target relationship was more predictive (*R* = −.035, *P* = .086 by bootstrap across participants). Formal analysis of optimal responding in two-choice reaction time tasks has shown that, normatively, uninformative cues should discourage deliberation, and lead to faster responding [24]. Consistent with this prediction, the speeding effect was significant when including only informative cues (60% and higher; *R* = −.050, *P* = .008). Finally, the longer participants had available to sample memory evidence, the more they responded early – early responses increased with ISI – but only when participants were signaled that the upcoming sensory evidence would be of low quality (low coherence: *R* = .110, *P* = .047; high coherence: *R* = −.003, *P* = .511; difference: *d* = 1.134).

Separately, the model predicts that responses made after the onset of the flickering probe stimulus should also reflect the quality of both kinds of evidence and, critically, now that both types of evidence are explicitly available, that these effects should interact with whether or not the content of memory and the probe are in agreement. Consistent with this model, for responses after the onset of the flickering stream, RTs were faster when the cue was more predictive (*R* = −.017, *P* = .044), and when the flickering stream was higher coherence (mean RTs – probe-locked, log-transformed, Z-scored within participant: low coherence: 0.158 SEM 0.061 high coherence: −0.128 SEM 0.039 mean difference between low and high coherence RTs within-participant 0.286, SEM 0.091, *t*(29) = 3.145, *P* = .004). These factors indeed interacted: participants were more speeded by cue predictiveness when coherence was lower (low coherence: *R* = −.147, *P* < .001; high coherence: *R* = −.065, *P* = .033; difference: *d* = 1.675), and only when the target photograph matched the cue’s prediction (valid cue: *R* = −.063, *P* < .001; invalid cue: *R* = .042, *P* = .141; difference: *d* = 1.739; Figure 3b).

Taken together, these results confirm that participants’ responses reflected the integration of information from both mnemonic cues and sensory input.

### Model comparison

We hypothesized that integration of mnemonic and sensory information resulted from dynamic, online inference on the basis of the quality of each source of evidence. We used formal model comparison to test this hypothesis.

Our primary model of interest implemented a continuous, two-stage sequential sampling process (hereafter: MSDDM; [21]; Figure 2b), – the first stage driven by the cue, and preceding the flickering probe, and the second stage beginning at the onset of the flickering stream – with different sampling rates in each stage.

An MSDDM is distinguished from alternatives by two key features: first, that the drift rate changes at the time of flickering stream onset, and second, that sampling in the second stage proceeds from the evidence sampled during the first stage. Therefore, we compared the model against variants that selectively disabled each of those features. The first comparison model was a single DDM, which had continuous sampling until the time of response, but no change in drift rate across the entire trial – i.e. responses reflected all available evidence up to that point, weighted equally regardless of modality. We refer to this model as *1DDM*. The second comparison model was two unconnected DDMs, mirroring the change in drift rate found in MSDDM, but with the second-stage starting point set independently of the behavior of the first stage – i.e. evidence sampled in the first stage only affected responses made before the onset of the flickering probe. We refer to this model as *2DDM*. Each model was fit separately to responses aggregated, across participants, by condition – cue, coherence, and ISI.

Against both comparison models, the MSDDM was a superior explanation of choices and response times. Against the second-best model – the 2DDM model – MSDDM was superior by BIC (BIC(MSDDM)= 1745.262, BIC(2DDM)= 10160.319, mean difference, across conditions: 187.00). This was the case across all conditions, and for every condition individually (Figure 4a; S1);

**Figure 4.**
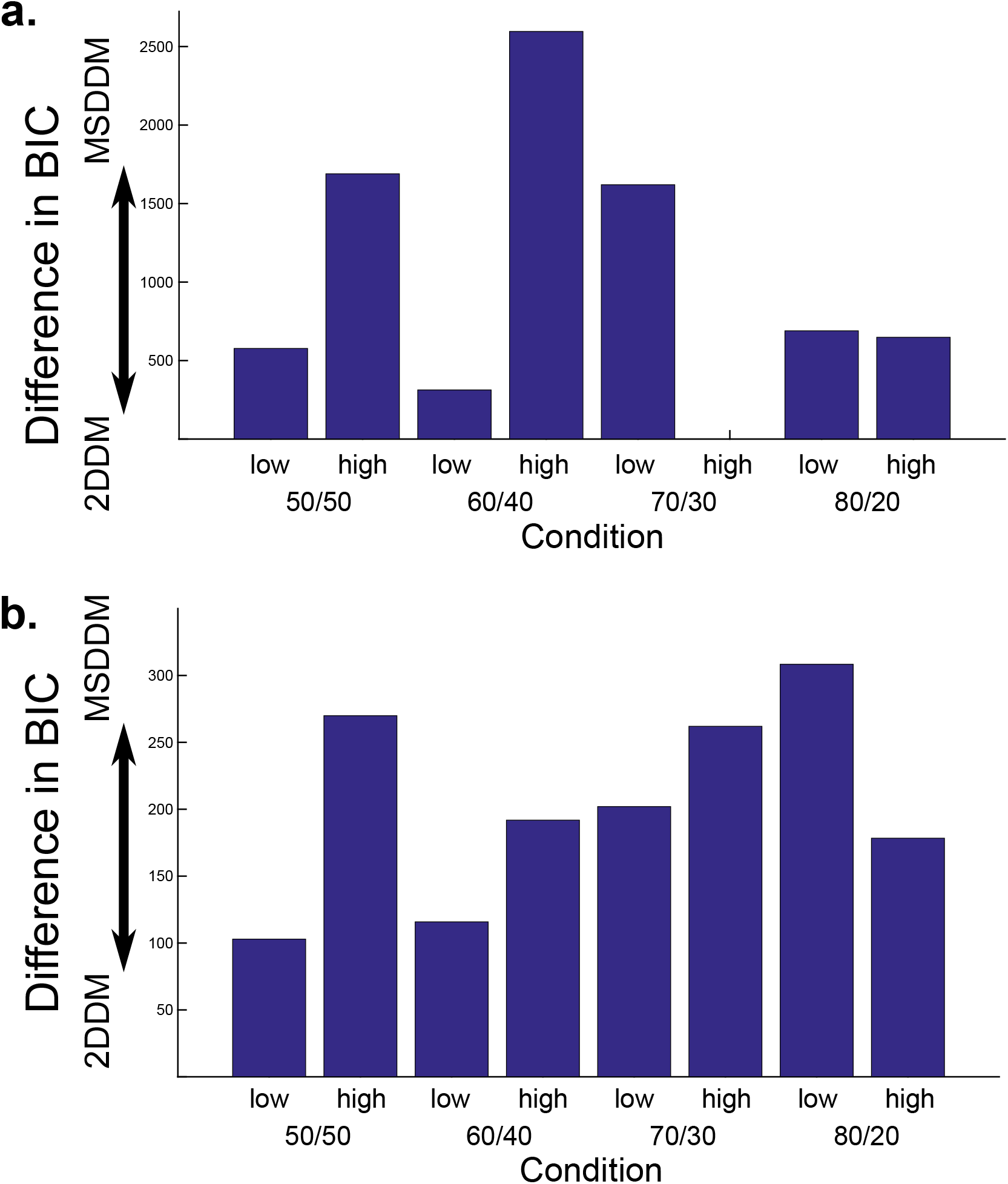
Model comparison. Models were compared for their fit to each bin of trials with the same combination of cue predictiveness, stimulus coherence, and ISI. Shown here is the BIC difference in favor of the MSDDM model (axes oriented such that a taller bar = greater evidence in favor of MSDDM), for conditions grouped by cue predictiveness and coherence level (aggregating across ISI, for display). In both experiments, MSDDM was favored over 2DDM for every condition individually, and across all conditions as a whole. **(a) Experiment 1**. BIC(MSDDM)= 1745.262, BIC(2DDM)= 10160.319. Mean difference, across conditions: 187.00. **(b) Experiment 2**. BIC(MSDDM)= 1230.385, BIC(2DDM)= 2861.661. Mean difference, across conditions: 67.97.

### Experiment 2

Experiment 1 showed that behavior in the task reflects a dynamic integration of memory and sensory evidence, yielding patterns of choices and response times that are best captured by the MSDDM. However, because they measure only the final response, behavioral data cannot in principle reveal a relationship between the actual memory evidence sampled on each trial and responses made to the ensuing flickering probe. In Experiment 2, we used multivariate pattern analyses (MVPA) of fMRI data to measure memory evidence samples following the fractal cue on each trial, and used this measure to predict responses after the onset of the flickering probe on that same trial. For this experiment, 31 additional participants completed the task from Experiment 1, while being scanned.

### Behavior

#### Response times and accuracy

Response behavior replicated the patterns observed in Experiment 1. Accuracy was again high overall: 70.24% correct responses (SEM 1.18%); and reliably above-chance for 49/52 blocks individually (all *p ≤*.073).

Accuracy again increased with cue predictiveness (*R* = .247, *P* = .005) and coherence (*t*(26) = −4.301, *P* < .001). RTs were again bimodal (all HDS*≥*0.102, all *P* < .001). Higher cue predictiveness resulted in a greater tendency to respond early (*R* = .149, *P* = .012), though, in contrast to experiment 1, the effect was specific to high-coherence trials (low coherence: *R* = −.102, *P* = .111; high coherence: *R* = .419, *P* < .001; difference: *d* = 4.213), perhaps reflecting that, for this group of participants, the rate of early responding was already at or near ceiling when participants anticipated low coherence stimuli. These early responses were faster when cue predictiveness was higher (*R* = −.077, *P* < .001); this was equally true at either coherence level (low coherence: *R* = −.231, *P* < .001; high coherence: *R* = −.229, *P* < .001; difference: *d* = 0.161).

Responses after onset of the flickering probe were again speeded by coherence (low: 0.227 SEM 0.042; high: −0.098 SEM 0.078; mean difference 0.325 SEM 0.098; *t*(30) = 3.326, *P* = .002), and by cue predictiveness, in both coherence conditions (low: *R* = −.092, *P* = .021; high: *R* = −.060, *P* = .013), moreso when coherence was lower (difference: *d* = 0.760), and when the target photograph matched the cue’s prediction: (invalid cue: *R* = .025, *P* = .303; valid cue: *R* = −.017, *P* = .191; difference: *d* = 0.945).

Finally, model comparison again favored the MSDDM over the alternative model (BIC(MSDDM)= 1230.385, BIC(2DDM)= 2861.661, mean difference, across conditions: 67.97; Figure 4b; S1; fitted parameters in Supplemental Tables S3,S4).

#### Neuroimaging

We used neural pattern similarity to measure the trial-by-trial influence of memory sampling on responses. For each participant, we localized regions in the ventral visual stream that were more active for face versus scene processing (FFA; [25]) and for scene versus face processing (PPA; [26]) (Figure 5a). We next computed activity patterns corresponding to each photograph, in the appropriate category-preferring region (faces in FFA, scenes in PPA). We refer to these picture-specific patterns as the *target patterns*. The target patterns were defined on the basis of data from an earlier response-training phase of the task, in which participants learned which keys were mapped to each picture (see *Methods*). Critically, because this response-training phase preceded the introduction of the fractal cues, these neural activity patterns were decoupled from the fractal cues that were later learned to predict the corresponding photographs.

**Figure 5.**
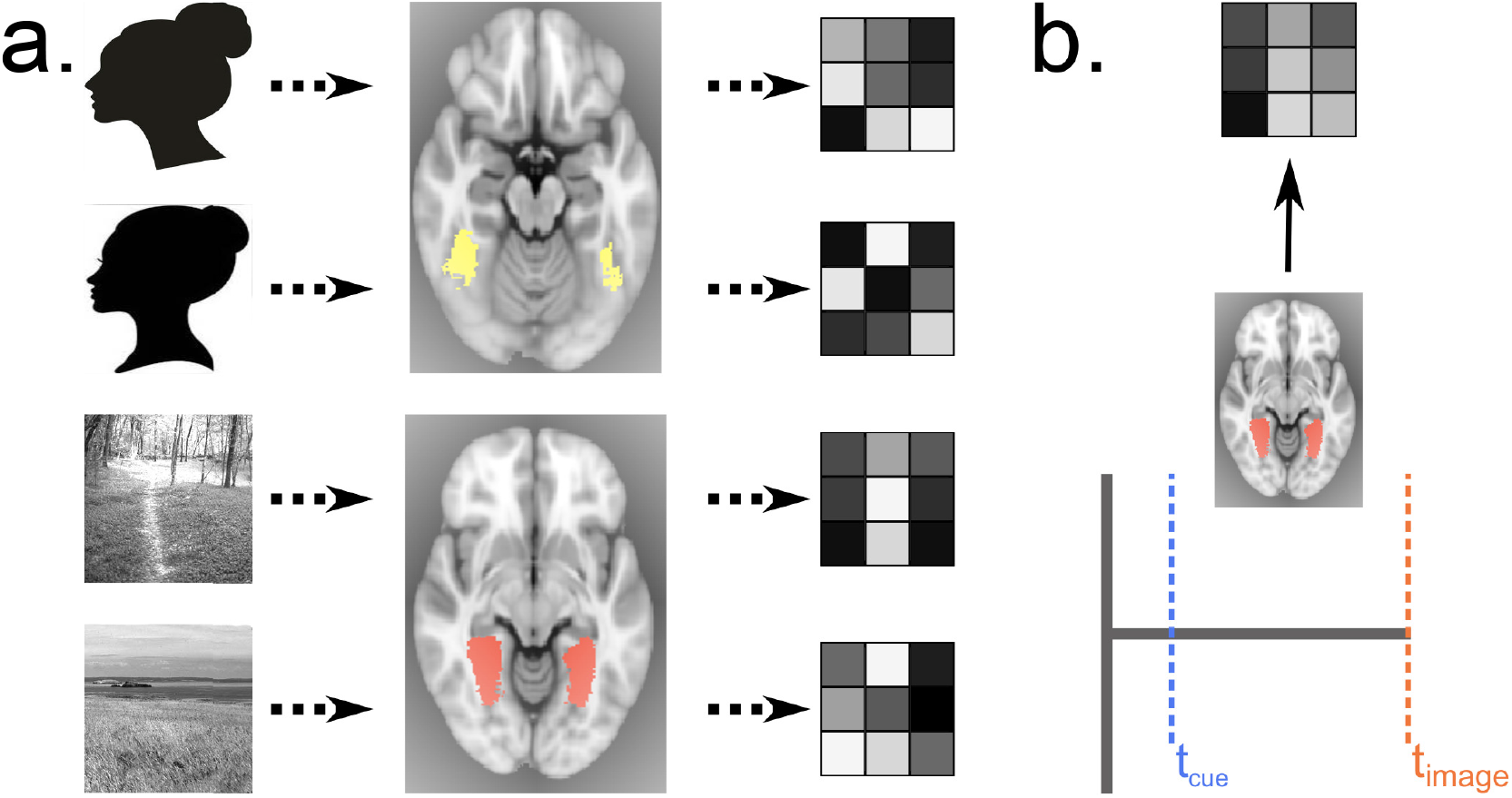
Reinstatement pattern analysis. **(a)** We defined, for each participant, the Fusiform Face Area (FFA) and Parahippocampal Place Area (PPA), using a localizer task that followed the main experiment. The resulting mask defined the region across which we calculated picture-specific templates. **(b)** For each trial on which participants responded after the onset of the flickering stream, we computed the average pattern of activity in the corresponding ROI (face, scene) over the period following the onset of the fractal cue, but preceding the onset of that flickering stream. We then calculated a *reinstatement index* as the correlation between this trial-specific pattern and the template pattern for the picture predicted by the fractal cue.

We then computed, for each trial from the Test phase, a *trial pattern* – the average pattern in these regions over the period following the onset of the fractal cue, up to either the participant’s response, or one TR before the onset of the flickering stream, whichever came first. Hereafter, we define the trial-by-trial *reinstatement index* as the correlation between these trial patterns and the target pattern corresponding to the photograph predicted by the fractal cue. (Note that on 50/50 trials this value is not defined, and so these trials were excluded from neuroimaging analysis.)

#### Pre-stimulus reinstatement scales with task conditions

As in a previous study of memory sampling [13], we expected that, when the next-step association was more difficult to resolve – when the cue was followed by each photograph more equally, in the participant’s experience – sampling should proceed longer, and thus more memory samples should be drawn across the anticipation period, leading to a higher value of the reinstatement index. Conversely, when sampling from memory reached thresh-old – and a response was initiated – there should be fewer reinstatements, and thus a lower reinstatement index. Similarly, memory sampling should continue across the entire anticipation period as long as participants do not respond before the flickering probe, leading to a higher reinstatement index with a longer anticipation period. Further, matching the patterns in early response times, memory sampling should be more relied upon when it would be more useful to the decision – in other words, reinstatement index should be higher when the upcoming sensory evidence would be of lower coherence.

The reinstatement index measure exhibited all of these features, consistent with the hypothesis that it measures memory sampling. On early response trials, reinstatement index was lower when the memory-based decision was easier (correlation between cue probability and reinstatement index *R* = −.072, *P* = .004; Figure 6a). On late response trials, reinstatement index was uniformly higher than on early response trials, and was equally high at every cue probability level (*R* = .020, *P* = .225; Figure 6a). Finally, reinstatement index was higher on trials with a longer ISI preceding low-coherence, but not high-coherence, stimuli (low coherence: *R* = .183, *P* = .016; high coherence: *R* = −.083, *P* = .247; difference: *d* = 1.546; Figure 6b).

**Figure 6.**
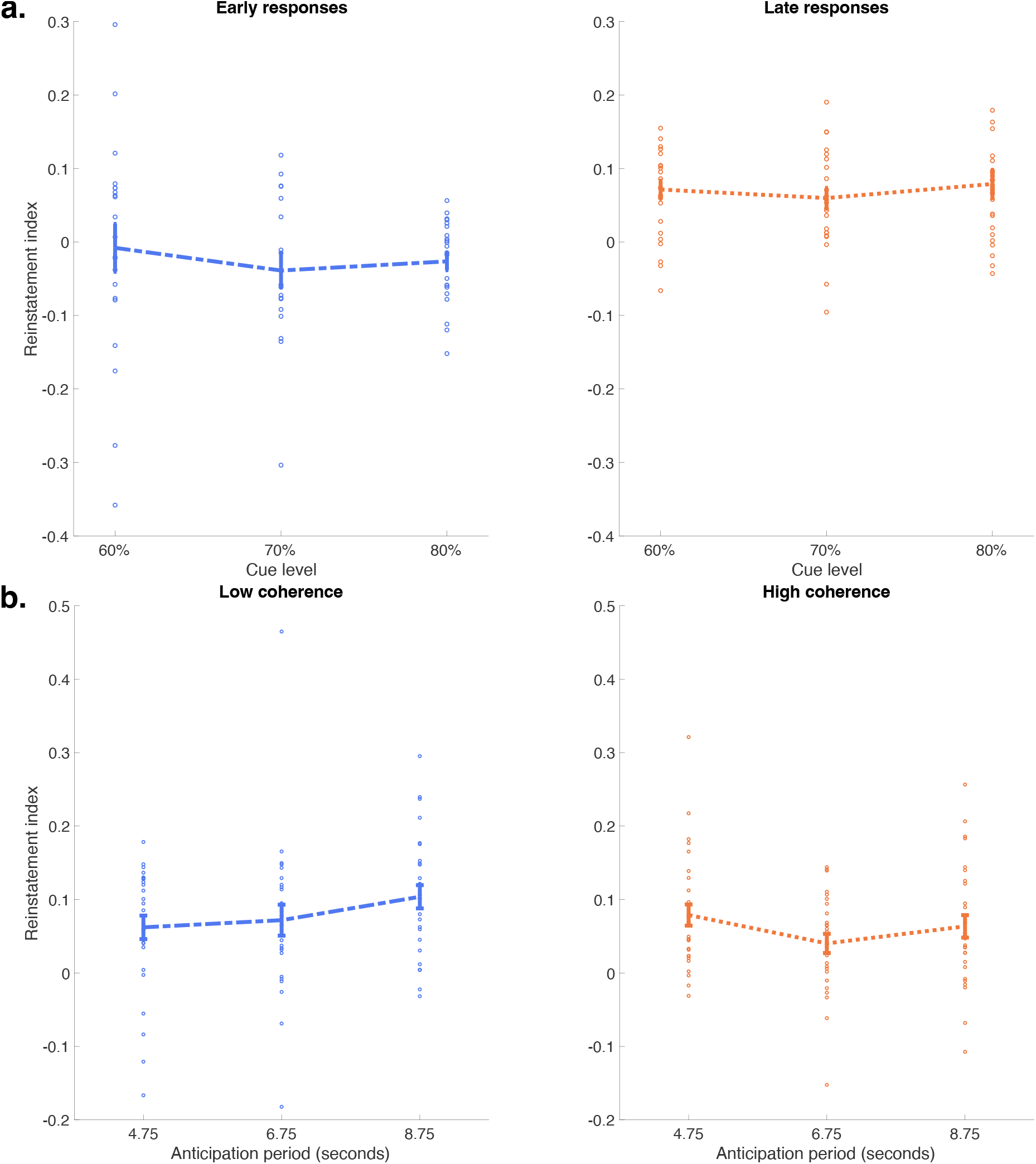
Reinstatement index measures adaptive sampling from memory. Reinstatement index changed with the quality of memory evidence, the anticipated quality of sensory evidence, and the available time to sample. **(a) Memory sampling increased with the difficulty of the memory decision**. When subjects responded “early” – in other words, when sampling terminated during the delay period – reinstatement index was higher when cues were less predictive of a unique target (lower cue probability; *R* = − .072, *P* = .004). When subjects responded “late” – by definition, sampling continuously throughout the delay period – reinstatement index was uniformly higher than on early response trials, in every condition individually, and across all conditions together (*t*(134) = −5.945, *P* < .001). Consistent with this measure indexing the subjective difficulty of the decision on each trial, the reinstatement index on late responses was uncorrelated with cue predictiveness (*R* = .020, *P* = .225). **(b) When sensory evidence was weak, memory sampling continued throughout the anticipation period**. On late response trials, reinstatement index increased with the length of the anticipation period – which allowed more time available to estimate the upcoming stimulus – but *only* on trials on which the upcoming sensory evidence was to be of low coherence (low coherence: *R* = .183, *P* = .016; high coherence: *R* = −.083, *P* = .247; difference: *d* = 1.546). (Error bars are *±* 1 SEM, across participants.)

The fact that reinstatement index is globally higher for late response trials, versus early responses (Figure 6a), raises the possibility that this measure is simply indexing the amount of data included in the analysis. Indeed, in the case of late responses, the reinstatement index is computed over the entire period between the cue and the visual evidence stream, whereas for early responses the measure only includes the time between cue and response (up to the limit of the temporal resolution of our fMRI protocol). An important validation that this difference in included timepoints is not driving the effect can be found in three features of the presented data. First, if the increase in reinstatement index were simply reflecting lower variability when including more data, then one would expect variance to decrease with longer response times during early-response trials. However, the variance of reinstatement index is in fact steadily greater for early responses to less-predictive cues, which are associated with longer response times (Figure 6a, left panel; 60%: SEM=0.032565, 70%: 0.023232, 80%: 0.01206). Second, while reinstatement index does indeed increase across the anticipation period for trials where the participant expects low-quality sensory evidence (Figure 66b, left panel), consistent with greater reinstatement activity during these longer anticipation intervals, the corresponding quantity remains flat or even numerically decreases across longer intervals for trials on which the participant expected high-quality sensory evidence (Figure 66b, right panel), consistent with the idea that anticipatory reinstatement on these trials proceeds at a slower rate (or perhaps stops early on). Finally, we observed that the variance of the reinstatement index neither decreases (nor increases) as a function of time alone, either across or within coherence conditions (Low-coherence: 4.75s: SEM=0.01602, 6.75s: 0.02107, 8.75s: 0.015815; High-coherence: 4.75s: SEM=0.014454, 6.75s: 0.013046, 8.75s: 0.015277).

#### Pre-stimulus reinstatement predicts 2nd stage response times

The observation that reinstatement index was related to cue probability is consistent with sequential sampling models, but does not itself differentiate between MSDDM versus 1DDM or 2DDM. Therefore, a key test of the MSDDM is whether responses to the flickering probe were influenced by sequential sampling in anticipation of sensory input, on a trial-by-trial basis. In other words, we asked whether memories retrieved *before* the probe affected responses made *after* its onset. To test this, we examined whether reinstatement index predicted response times after the onset of the flickering probe on each trial.

Supporting our hypothesis, reinstatement index was a reliable predictor of faster response times to the probe (*R* = −.07, *P* = .001), a relationship that held after controlling for other factors each of which also modulated response times (cue predictiveness, coherence, ISI; *R* = −.0337, *P* = .039). If sampled memory evidence sets the starting point for sensory inference, then reinstatement index should only predict faster responses when memory and sensory evidence are in agreement – when the cue is “valid.” In agreement with this hypothesis, RTs were uniquely speeded on valid- – but not invalid- – cue trials (valid cue: *R* = − .053, *P* =.029; invalid cue: *R* = .010, *P* = .404; difference: *d* = 1.204; Figure 7a). Finally, if memory and sensory evidence were integrated, memory evidence should show correspondingly less influence when sensory evidence was stronger; this should be reflected as a greater speeding of matching, relative to non-matching, trials. Consistent with this prediction, the benefit to memory evidence was pronounced in the low-coherence condition, and no such benefit was observed in the high-coherence condition (low coherence, invalid cue: *R* = .130, *P* = .087; low coherence, valid cue: *R* = −.063, *P* = .032; difference: *d* = 1.586; high coherence, invalid: *R* = −.089, *P* = .117; high coherence, valid: *R* = −.037, *P* = .206; difference: *d* = 0.423; Figure 7b).

**Figure 7.**
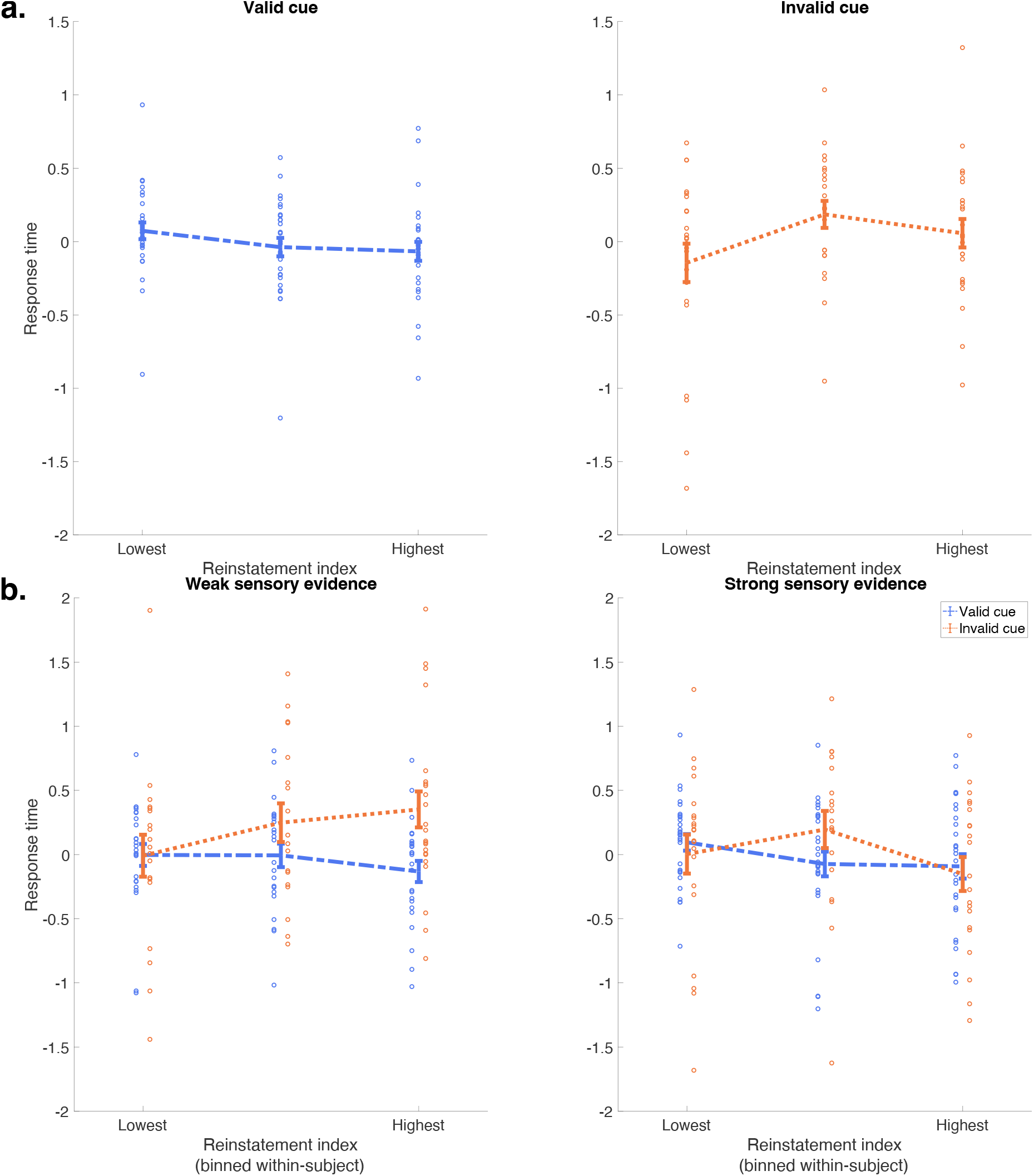
Sampled memory evidence is incorporated into perceptual decisions. (a) Memory evidence speeded responses when the target matched expectations. The MSDDM predicts that sampled memory evidence should carry forward to visual inference, effectively setting the starting point for this second stage. Thus, if the visual stimulus matches the sampled memory evidence, responses should be speeded, and, if the visual stimulus doesn’t match the sampled memory evidence, responses should be slowed. Critically, this speeding or slowing should be dynamic, corresponding to the strength of evidence sampled *on that trial*. Consistent with this model, higher reinstatement index predicted faster responding to sensory information when the cue was “valid”, or matched the sensory stimulus, but not when the cue was invalid (valid: *R* = −.053, *P* = .029, invalid: *R* = .010, *P* = .404, difference: *d* = 1.204). **(b) Memory evidence affected responses to weak, not strong, sensory evidence**. Further confirming the model, the speedup and slowdown was pronounced on trials where sensory evidence was weaker (low coherence: valid: *R* = −.063, *P* = .032, invalid: *R* = .130, *P* = .087, difference: *d* = 1.586), but less so when sensory evidence was strong (high coherence: valid: *R* = −.037, *P* = .206, invalid: *R* = −.089, *P* = .117, difference: *d* = 0.423). (Error bars are *±* 1 SEM, across participants.)

Together, these results support a role for the dynamic sampling of memory evidence in perceptual decisions, via a continuous perceptual inference process linking memory, sensation, and action.

## Discussion

Humans [6, 27], animals [11], and even intelligent machines [28] rely on expectations, derived from experience, to make quick and accurate decisions [24]. While important empirical and theoretical work has described ways in which expectations influence dynamic, deliberative decisions [6, 7], these investigations have generated seemingly-conflicting results [5, 7, 29, 30], and have mainly set aside the dynamics of expectation-setting itself. These studies generally assume that subjective expectations are more or less static across repeated trials of the same type [5, 11, 29, 31], and are instantaneously available to the subject at choice onset. These assumptions are justified when expectations can be based on extensive repeated experience in stationary environments, or explicit instructions. However, even despite designs that muted the potential for expectations to vary between instances of a decision, such variation is still commonly observed. A standard approach is to model this variance as random Gaussian fluctuations of the parameter values [32, 33]. It is an open question what mechanism gives rise to this variation.

We reasoned that, if trial-by-trial variability in expectations is a signature of a dynamic process, this variability would be most pronounced – and thus most subject to interrogation – when expectations were based on relatively few experiences, and when there was sufficient time available for these experiences to be sampled before the decision. Our study sought to characterize and explain trial variability in expectations by examining a space left unaddressed by previous work: that in which the dynamics of expectation setting could be separately manipulated, and measured, on a trial-by-trial basis.

Participants performed a novel cue-guided perceptual decision task, in which they were asked to press a key corresponding to what they believed to be the target photograph on a given trial. On each trial, prior to probe presentation, memory cues triggered reminders of past sequential associations between the cue and target or non-target photos. Then, after a delay, a probe stimulus displayed the target and non-target, in a noisy, rapidly flickering stream. Thus, both parts of the trial provided partial information as to the correct response. Choices and reaction times reflected the combination of both kinds of information, and were best fit by integrative two-stage sequential sampling of memory and sensory evidence. Using fMRI, we showed that second-stage responses on each trial were predicted by the reinstatement of stimulus-specific representations, the content and fidelity of which fluctuated between trials, that developed with the time available to sample on each trial, and the content of which reliably predicted the pattern of behavioral response. The results demonstrate that sensory evidence sampling in perceptual decisions reflects the influence of a preceding phase of an inference process that is already ongoing at the onset of the visual probe.

Several recent findings can be reinterpreted in light of a continuous inference mechanism that begins before the onset of sensory information. Multiple reports have observed, using electrophysiology in non-human primates, “ramping” of neural activity in putative accumulator regions during epochs preceding sensory and motor decisions [11, 34]. Hauser and colleagues (2018) observed that pre-trial ramping activity had a meaningful influence in explaining RT variability. Specifically, they identified ramping with variability in the onset of motor plans. They ascribed this to a mechanism that considers countermanding motor plans. Consistent with this view, in our model fits the second-stage non-decision time showed a trend towards decreasing with increasing cue predictiveness (Supplemental Figure S3) – in other words, less onset delay when discordant outcomes are less likely to be sampled. From this perspective, our model suggests that the onset variability they identified reflects the degree to which preparatory activity, reflecting inference about trial expectations, conflicted with sensory information, thus delaying action.

Hanks and colleagues (2011) reported a small amount of pre-trial ramping, but found that it did not have direct impact on subsequent behavior. It seems likely that two features of their design worked against such a result. The first is a feature that, as we noted above, is common to most studies involving expectation-setting: fixed priors learned to asymptote over many hundreds of experiences. The second feature is the use of short inter-trial-intervals, on the order of 100s of milliseconds, potentially insufficient to permit the development of memory-guided expectations. While these aspects of their design were critical to the goals of their study, which focused on the equilibrium dynamics of expectations in inference, they are likely to diminish the need and/or opportunity for online inference informing the content of expectations. Indeed, a possible unification of our results and theirs is that the dynamic bias signal they observe is ongoing inference over the prior [35], and that the short anticipation delay meant that the expectation-setting process (proceeding in parallel to sensory inference; see below) only accumulated to an observable degree when sensory decisions persisted long enough. Informed by our previous work on memory-guided, value-based decision-making [13, 15, 16], the present study was designed with these considerations specifically in mind. Further work is necessary to understand the precise conditions under which expectation-inference does and does not have meaningful influence on the subsequent decision.

As we mentioned above, it has been observed that behavior in sensory inference tasks tends to be best fit when model parameters are allowed to vary – stochastically – between trials [32]. This is true both when the parameters are set on the basis of known task conditions (in other words, expectations) – for instance, when the drift rate is enhanced for a more common stimulus [36, 37], or starting point is set to favor the more common response [5] − and also when they are allowed to vary freely [33]. In both cases, such variability might arise from active sampling of recent past trials, the outcomes of which would bias sampling model parameters towards a faster, more accurate response in that previous condition. Intriguingly, in the latter case, this would imply that inference is ongoing even when long-run expectations are not justified [38]. It might thus be worth investigating whether signatures of a pre-trial inference process – e.g. lower variance of diffusion parameters following longer inter-trial-intervals (Supplemental Figure S2) – are present even in tasks without decisive expectations.

The use of memory samples to set inferential decision parameters in this study is also motivated by work on reinforcement learning, in which memory-based planning is used to accelerate learning by reducing uncertainty in the agent’s policy and state representation. It has been shown that such memory-guided planning is particularly useful within complex or “partially observable” state spaces [39, 40], and early on in learning of even simple tasks [41]. We and others have also recently shown that the influence of memory on decisions can be specifically biased by reminder cues presented before the onset of choice options [15, 16, 42, 43], similar to the informative fractal cues presented here. In light of the present findings, a unifying explanation of these results is that the brain is engaged in an ongoing attempt to reduce uncertainty about the current state by drawing on memories of similar past situations and that such reminder cues, even when incidental, are treated as meaningful indicators of similarity.

This interpretation suggests one answer to the question of why participants in this task use memory samples – even though the task is designed in such a way that memory samples cannot, on their own, be decisive. Returning to the reinforcement learning terminology, we can understand the sensory decision here as one instance of a broader class of problems, those of *state inference*. Though the task used here is modeled most directly on the canonical perceptual decision process that between two actions on the basis of a noisy stream of visual information, Rao (2010) has shown that such sequential sampling models like we use are formally equivalent to a rational approach taken by a reinforcement learning agent navigating a partially observable markov decision problem (POMDP). In a POMDP, as opposed to the fully observable MDP, the agent acquires only partial information about the current state − e.g. the noisy sensory information and, in our task, the history of experiences with the state that followed the given fractal cue. Their internal state representation is probabilistic − a distribution of “beliefs” over the current state, and action selection is thus a process of jointly inferring both the state and the action contingencies that follow. This distribution reflects both the estimate of the current state and the uncertainty the agent has about that estimate. An agent navigating this environment has three actions available to it – the two motor responses, and a third “sample” action, in which it chooses to acquire additional evidence that may help reduce uncertainty in its belief distribution, rather than to act externally. Because new evidence from memory can do no worse than increase certainty in the current action policy [40], the expected value of acquiring new evidence samples is strictly non-negative, net the cost of acquiring a sample (e.g. a metabolic cost of memory retrieval, a foregone reward in delaying action, or foregoing the opportunity for a few seconds of rest). The agent therefore has incentive to continue taking that “sample” action as long as it is useful for reducing uncertainty, over and above any costs. The use of memory samples implies that the cost of acquiring a memory sample must be less than the initial value of reducing this uncertainty in this task.

A related question is why participants respond early, before the onset of sensory information. A similar finding was reported by Kiani and colleagues [44], who observed that monkeys performing a direction-discrimination task of experimenter-controlled duration committed to a decision once accumulated evidence crossed the monkeys own internally-set threshold, regardless of whether more, potentially decisive, evidence was yet to come. The authors of that study interpreted their finding as potentially indicating a “cost” of sampling or, equivalently in both their and our setting, a cost to waiting to act, unrelated to the timing of the task itself (their task, like ours, was of fixed length and thus did not reinforce faster responding). What exactly gives rise to this cost remains an open question. Further work is necessary to understand the internal trade-offs made when weighing the costs and benefits of sampling from memory and the environment.

The finding that multiple streams of evidence are integrated raises the question of whether such integration is purely serial, or if it can operate in parallel. In other words, does memory sampling cease to influence choice at the time of the onset of the flickering probe – or even before (e.g. “freezing”; Kiani et al. 45, Hoskin et al. 46) – or does it continue once that imperative stimulus has appeared, contributing additional evidence from expectations even as the sensory decision unfolds [11]? Our model is consistent with both possibilities, because the second-stage drift rate that we fit can be equivalently interpreted as either the drift rate of sensory accumulation or of the superimposed sampling from both memory and sensory evidence. Addressing this question further will likely require neural recordings at fine temporal resolution and broad spatial coverage, in order to identify separate, simultaneous sampling timeseries.

Whether sampled in serial or parallel, each form of evidence must be weighted in its contribution to the final action selection. What is the proper weighting of each, and how is it determined? Here we show that the two kinds of information are mixed, and that the weighting is determined in part by the dynamics of sampling each representation within each trial, and also, implicitly, by an exogenous mechanism that decides how much to sample from memory in the first place. Is this decision made at the cue presentation? Or is it also a dynamic decision – are samples first drawn and then evaluated for their informational content? Our ability to resolve these questions in this experiment was limited by the temporal resolution of our measurements as well as aspects of the design: in this experiment the sensory information was – inherently – more decisive than the memory. A follow-up experiment, perhaps reversing the order of presentation of memory cues and flickering probe [47], could build on the results and tools demonstrated here to more finely measure the effective weight in behavior, and how that weight is determined at the level of neural activity.

One such determining factor might be the degree of confidence in the initial estimates. Previous work has shown that the brain codes measures of confidence suitable for determining weighting of the sensory evidence necessary to select action [48–50], and that this measure predicts subsequent “changes of mind” on the basis of late-arriving sensory evidence [51, 52]. Recent work also suggests that a corresponding quantity is computed separately for memory evidence [53, 54]. Do these confidence estimates inform the mixture of multiple kinds of evidence in behavior? A related question is whether, across modalities, such measures are absolute, reflecting only the confidence within the given representation, or are they coded relative to the confidence available in other representations? By setting the parameter selection exogenous to this model, we also set exogenous the likely mechanism for confidence to influence the process. It is possible that the degree of confidence in evidence available on the first stage can inform the parameters of the second stage – and vice-versa, since the quality of sensory evidence is signaled ahead of time [55, 56]. Future extensions of this task should bring such decision-relevant confidence under experimental measurement or control.

Along similar lines, recent work has shown that multiple forms of associative maps can be learned and represented simultaneously, reflecting both statistical and semantic relationships among stimuli [57]. This finding raises the question of whether these multiple maps also simultaneously contribute to online inference of the sort examined here, a possibility formalized in a recent theoretical framework describing the dynamics of parallel sampling from internal representations [56]. According to this framework, each set of associations should guide behavior in proportion to its relative precision at each moment [55]. However, as described above, the current experiment approach was limited in its ability to discern whether the multiple forms of evidence are integrated in parallel. Future studies could select cue-target associations that have both in-task statistical and, e.g., semantic relations, and examine factors that modulate the relative influence of both kinds of information on behavior.

A final caveat is that our results do not speak to the question of how evidence samples are translated into decisions. A groundbreaking recent computational investigation recently demonstrated that the reaction time properties consistent with the entire class of evidence accumulation models, such as the one we fit to data here, are also consistent with a different set of approaches that, rather than accumulating evidence across multiple samples, simply select an extreme or most-relevant individual sample [58]. In light of this important finding, it is necessary to make clear that our study, like nearly every other investigation of evidence-sampling behavior in recent decades, cannot in principle directly determine the algorithmic-level process underlying the expectation-setting process (e.g. accumulation versus extrema detection). We anticipate that future studies, in particular those which employ methods using high temporal resolution paired with broad spatial coverage, may be able to integrate our observations with the results of Stine and colleagues to address this foundational question. To more directly address the accumulation question, future studies, perhaps employing measurement protocols with wider coverage, could also examine whether the re-instatement index predicts more general accumulation activity in downstream regions (e.g. LIP or anterior frontal areas). Such studies could examine in more detail our observation that reinstatement index (measured during the cue period) is sensitive to upcoming visual coherence – suggesting that sampling may be minimal, and perhaps restricted to early time-points, during the anticipation of high-coherence trials, reflecting the relative likelihood of high-quality visual evidence soon to appear.

Because expectations are nearly omnipresent in decision-making, it is possible that previous investigations have obscured an important source of trial-by-trial variation. Decisions may often be biased by samples from internal information – memories, but also emotions, values, and rules – that give rise to expectations established in the moment, rather than fixed across time. Biases, derived from experience, are helpful, and under some circumstances, may even be necessary, for efficient decision-making – they help us take account of, and leverage, the statistics of our environment. Current research is outlining a role for goal-directed, decision-time planning in a number of psychiatric conditions. Dysfunction in this mechanism could explain disease states characterized by under-or over-reliance on expectations in behavior such as in Parkinsonism [9, 10], disorders of compulsion [59], or positive symptoms in schizophrenia [60]. Consistent with the observation of Perugini and Basso (2017) that deficits in expectation-setting cannot be explained by altered dopamine function, the present findings provide an aperture for treatments, by underscoring that biases are not simply “stamped-in” regularities. Instead, the fact that expectations are constructed in the moment implies that they can be changed, via targeted interactions with the construction process. Outside of disease, biases alterable in the service of better decisions may be a crucial adaptation, allowing organisms to adjust their behavior at a timescale faster than that of the long-run statistics of their environment. More broadly, it means that, when it comes to individual decisions, the link between past and present can be revisited, even changed, when the need arises.

## Acknowledgements

The authors wish to thank Abigail Hoskin, Amitai Shenhav, Judith Fan, Phillip Holmes, Michael Waksom and Roozbeh Kiani for helpful conversations, Ghootae Kim for providing ranked face and scene stimuli, Nicholas Hindy for providing fractal stimuli, and Charlotte Townsend for extensive assistance with data collection. This publication was made possible through the support of funding from the Intel corporation, and a grant from the John Templeton Foundation (Grant ID #57876; K.A.N. and J.D.C.). The opinions expressed in this publication are those of the authors and do not necessarily reflect the views of the John Templeton Foundation.

## Open practices statement

The data and materials for all experiments are available at OpenNeuro (doi:10.18112/openneuro.ds001614.v1.0.1). Neither of the experiments was pre-registered.

## Contributions

A.M.B. and M.A. conceived experiment; A.M.B., M.A., and S.F.F. designed experiment and analyses, with input from N.B.T., K.A.N. and J.D.C.. A.M.B. and M.A. wrote the experiment code; A.M.B. and M.A. ran the experiment; A.M.B. and S.F.F. contributed analytic tools; A.M.B. and S.F.F. performed analyses; A.M.B. wrote the paper, with input from M.A., N.B.T., K.A.N., and J.D.C.. All authors approved the final manuscript.

## Methods

### Participants

33 participants (15 male, 30 right-handed; ages 18-50, mean 21.9) each performed two repetitions of the task in Experiment 1. Ten blocks were excluded for failing to meet one or more criteria: if the participant failed to respond on 10% of learn or test-phase trials (nine blocks); if the combined number of skipped trials and post-stimulus error trials during the test phase were greater than 30% (four blocks); if the difference between calibrated accuracies for any pair of stimuli was less than 5% (one block). Three participants failed to meet criteria for all blocks they performed; they were excluded entirely from analysis. In all, 30 participants and 56 blocks were included in the final analysis.

36 participants (10 male, 29 right-handed; ages 18-33, mean 23.19) each performed one (5) or two (31) repetitions of the task in Experiment 2. (Five blocks were excluded due to scanner malfunction (1), participant discomfort (1), or programming error (3).) 15 blocks were excluded for failing to meet one or more criteria: nine for failing to respond on enough learn or test-phase trials; one for failing to respond correctly or at all on enough test-phase trials; nine for failing the calibration accuracy threshold. Five participants failed to meet criteria for all blocks they performed; they were excluded entirely from analysis. In all, 31 participants and 52 blocks were included in the final analysis.

In Experiment 1, participants were compensated with course credit. In Experiment 2, participants were paid a flat fee of $50. All participants reported themselves as free of neurological or psychiatric disease, and fully consented to participate. The study protocol was approved by the Institutional Review Board for Human Subjects at Princeton University.

### Task

The experiment was controlled by a script written in Matlab (Mathworks, Natick, MA, USA), using the Psychophysics Toolbox [61]. Both Experiment 1 and Experiment 2 consisted of the following four phases, repeated for two blocks for each participant, with different stimuli and task conditions as detailed below. Experiment 2 was performed in an fMRI scanner, and consisted of an additional, fifth phase, a Localizer task described below.

In *Phase 1*, the *Response training* phase, particpants learned to map response keys to stimuli. Four response keys – numbers one through four on a standard US keyboard – were each associated with one of four stimuli – black and white photographs, two faces and two natural scenes.

Stimulus photographs were chosen from a set of four possible scenes and four possible faces. Each category was subdivided into two sets of two paired photographs. Each photograph was black and white, normalized for contrast and brightness, and chosen to be highly confusible with the paired face or scene.

Participants were first shown each photograph, centered on a black background, in order of the associated response keys, and asked to press the current key in the sequence. In all experiments. keys one and two corresponded to the faces, and keys three and four corresponded to scenes. Then, the photographs were shuffled, and presented one at a time for two seconds each. Participants were instructed to press the corresponding key. If they pressed the correct key, a green box appeared around the photograph. If they pressed the incorrect key, the photograph remained on the screen. Each photograph was displayed ten times. If participants pressed the incorrect key on the first try more than twice for any photograph, they were made to repeat the response training phase in its entirety.

*Phase 2*, the *Calibration* phase, measured the ability of participants to discriminate between each pair of photographs when they were presented in a noisy, “flickering” stream (Figure 1). On each trial, participants were shown a rapid stream of pictures, displayed for 1/60th of a second apiece. They were instructed to press the key corresponding to the *target* − the photograph shown most often. Each frame consisted of either the target photograph, the paired same-category photograph, or a perceptual mask consisting of a phase-scrambled version of a superposition of the two photographs. Perceptual masks were shown for between one and three frames, with mask display length chosen from a truncated, discretized exponential distribution of mean 2. Calibration trials lasted three seconds, regardless of response. When participants pressed a key, the stream stopped, and the target was shown for the remainder of the trial length. If the participant pressed the correct key, a green box appeared around the photograph. If the participant pressed the incorrect key, a red box appeared around the photograph. A one second inter-trial-interval (ITI) followed each trial. On each trial, the proportion of frames that contained the target photograph – the *coherence* − was updated according to a Quest algorithm [62], with the goal of calibrating participants responses to either 65% (low) or 85% (high) accuracy. Each block measured the coherence necessary to elicit either high or low accuracy for each photograph. In Experiment 1, the first 24 participants performed 60 calibration trials per photograph, while the last 9 participants performed 40 calibration trials per photograph. In Experiment 2, participants performed 30 calibration trials per photograph. Although Experiment 2 participants remained in the fMRI scanner for this phase, no scanner data was collected. This is the only phase for which scanner data was not collected.

In Experiment 1, for stimuli calibrated to low accuracy (65%), the average coherence (proportion of non-mask frames that contained the target photograph), across participants, blocks, and stimuli, was 60.98% (SEM 1.06%); whereas for the high-accuracy (85%) condition, the target photograph was shown on 75.88% (SEM 1.08%) of frames. In Experiment 2, these figures were 62.17% (SEM 1.03%) coherence in the low-accuracy condition, 77.66% (SEM 1.15%) coherence in the high-accuracy condition.

*Phase 3*, the *Sequence learning* phase, provided participants with a set of experiences that linked each of four fractal cues to the photographs (Figure 1). On each trial, participants were shown one of four fractal cues, displayed on the screen for 750ms. Fractal cues were chosen in order to minimize the possibility of pre-existing relationships between the cue and target, and thus to allow us to more directly examine the influence of statistical learning from a small number of experiences on subsequent perceptual decisions. In Experiment 1, the cue was followed by a variable inter-stimulus-interval (ISI). For 24 participants, this ISI was either 500ms, 1000ms, or 4000 ms, selected pseudorandomly at each trial according to a uniform distribution. For the remaining 9 participants in Experiment 1, and all participants in Experiment 2, this ISI was a fixed length of one second. After the ISI, participants were shown either of two photographs linked to the cue, both from the same category (face or scene). The photographs that followed the cue were selected according to one of four binomial distributions – 50/50, 60/40, 70/30, or 80/20. The two cues in each category (face or scene) predicted their consequents using symmetric distributions – if one cue predicted Face A with 80% probability, the other cue predicted Face B with 80% probability. Participants were instructed to press the button corresponding to the displayed photograph. If the response was accurate, the photograph was surrounded by a green box. If the response was inaccurate, the photograph was surrounded by a red box. Regardless of response time or accuracy, the picture remained on the screen for two seconds. In Experiment 1, the trial was followed by an ITI of two seconds. In Experiment 2, the trial was followed by an ITI of between 500ms and 8000ms, chosen from a truncated exponential distribution, discretized in units of 500ms, with mean 2000ms. This phase consisted of 100 trials, 25 for each cue, ordered pseudorandomly.

*Phase 4*, the *Cued inference task* was the primary test of our hypotheses. On each trial during this phase, participants first viewed a fractal cue that predicted the likelihood of the target photograph during the following flickering stream. Cues were presented for 750ms, and followed by an ISI of variable length, selected at each trial from a uniform distribution. For the first 24 participants of Experiment 1, this ISI was either 500ms, 1000ms, or 4000ms. For the remaining 9 participants of Experiment 1, this ISI was either six, eight, or ten seconds. In Experiment 2, this ISI was either four, six, or eight seconds. In both experiments, ISI durations were chosen from a uniform distribution over the possible values. The flickering stream used one of the two mixture proportions calibrated during Phase 2; mixture proportions were fixed for each category – e.g. faces might be set to low coherence, and scenes to high coherence. Thus, the fractal cue predicted both the likely identity of the target photograph, and also the coherence of the subsequent stream. The stream remained on the screen for three seconds. When a key was pressed, the target photograph appeared, and remained on the screen until the three seconds were finished. If the keypress was correct, the photograph was surrounded by a green box. If the keypress was incorrect, the photograph was surrounded by a red box. Participants were instructed to press the key corresponding to the identity of the target photograph. Critically, however, participants were allowed to respond early – during the ISI, before the flickering stream began. Participants were not given any explicit or implicit inducement to respond early or accurately – they were informed that, regardless of the speed or correctness of their response, all trials were of fixed length, modulo the ISI. This phase continued for 80 trials, 20 trials of each cue, ordered pseudorandomly.

Phases one through four were repeated as two blocks, each with different fractal cues and picture stimuli. Cue were selected pseudorandomly for each block, and the mapping from coherence level to category was counterbalanced between blocks.

After the two blocks, Experiment 2 participants completed a final phase, *Phase 5*, the *Localizer* task. We used the data collected in this phase to localize regions of cortex preferentially active during processing of face and scene images. Participants performed a 1-back image repeat detection task. Images were presented in mini-blocks of 10 trials each. Eight of the pictures in each block were trial-unique, and two were repeats of the picture on the immediately preceding trial. Repeats were inserted pseudorandomly according to a uniform distribution. Stimuli in each mini-block were chosen from a large stimulus set of pictures not used in the main experiment. The pictures belonged to one of four categories – faces, objects, scenes or phase-scrambled scenes. Pictures were each presented for 500ms, and separated by a 1.3s ISI. A total of 12 mini-blocks were presented (3 per category), with each mini-block separated by a 12 second inter-block interval.

### Imaging methods

Experiment 2 was collected while participants were laying in the fMRI scanner. Data were acquired using a 3T Siemens Prisma scanner with a 64-channel volume head coil. We collected three functional runs with a T2*-weighted gradient-echo multi-band echo-planar sequence (44 slices oriented parallel to the long axis of the hippocampus, 2.5mm isotropic resolution, echo time 26 ms; TR 1000 ms; flip angle 50 deg; field of view 192 mm). To allow for T1 equilibration, we discarded the first six volumes of each functional run (6s). We also collected a high-resolution 3D T1-weighted MPRAGE sequence (1mm isotropic resolution) for registration across participants to standard space. Functional image preprocessing was performed using FSL (FMRIB Software Library version 5.0.8; [63, 64]). Anatomical images were coregistered to the standard MNI152 template image, then individual participant functional images were coregistered to the realigned anatomical images. The transformation matrices generated during this coregistration process were used to transform Region of Interest (ROI) images (described below, *ROI definition*). Functional images were motion corrected and spatially smoothed using a 5mm full-width half-maximum Gaussian kernel prior to analysis. Data were scaled to their global mean intensity and high-pass filtered with a cutoff period of 128s.

### Behavioral analysis

#### Response time analyses

##### Bimodality

We tested whether response time distributions within each ISI condition were bimodal, using Hartington’s Dip Test [23]. This test measures the relative spread between modes to the mean of the distribution – larger values indicate a higher likelihood of true bimodality in the tested data. P-values are estimated via bootstrap against distributions with the same summary statistics as the tested data, provided by the MATLAB function H<sc>ARTIGANS</sc>D<sc>IP</sc>T<sc>EST</sc> [65].

##### Permutation tests for across-condition correlations

Each participant performed a different subset of the task conditions (cue level, perceptual coherence). To provide a robust measure of the relationship between response times and conditions, we therefore performed a bootstrap analysis, across participants and conditions [66]. On each iteration, we sampled, with replacement, the number of participants in the study group (30 in Experiment 1, 31 in Experiment 2). We then computed, on this selected group, the correlation of interest. By repeating the process 1,000 times, we obtained a distribution of correlation values across shuffled permutations of the study group. The reported p-value is thus the fraction of correlation values with a different sign from the base effect size (the correlation across the entire original group). When evaluating whether these correlations differed between conditions (e.g. for coherence levels, or for early versus late responses), we compared the difference between the values obtained for paired bootstrap iterations (using the same selected subset of subjects). For these tests, *P* -values that result from standard nonparametric tests are, generally, trivially significant, due to the large population size. Therefore, to evaluate the reliability of the difference we used Cohen’s *d* [67]; by convention effect sizes measured in this way greater than 0.80 are “Large”, and thus reliable.

#### Model comparison

##### Multi-stage DDM

Our primary model of interest is an extension of the drift-diffusion model [3]to allow for a time-varying drift rate [21]. The model specifies drift rate as a piecewise constant function, in which each shift in drift rate defines a separate “stage” of the sampling process. Critically, the endpoint of one stage naturally sets the starting point of the next. Our instantiation used two stages. The free parameters were thus the drift rates, *d*_1_ and *d*_2_, nondecision time *T*_0_, and a distribution of trial-by-trial first-stage starting points specified by the mean *x*_0_ and standard deviation 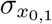 We refer to this model as *MSDDM*. Our comparison models were matched sequential sampling processes that each selectively disabled one key feature of the MSDDM – the time-varying drift rate, and the connection between stages. The first comparison model of interest was a single DDM, with continuous sampling until the time of response, but no change in drift rate across the entire period between the onset of the fractal stimulus and response. We refer to this model as *1DDM*, with free parameters *d*_1_, *T*_0_, *x*_0_, and 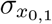. The second comparison model of interest was two DDMs, each fit to pre-stimulus and post-stimulus responses separately and thus mirroring the change in drift rate found in MS-DDM, but with the second starting point its own free parameter.

We refer to this model as *2DDM*, with free parameters *d*_1_, *d*_2_, *T*_0_, and starting points for each stage, defined by *x*_0,1_, 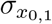 and *x*_0,2_, 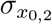. Each model was fit to participant responses aggregated according to cue, coherence, and ISI condition. The fitting procedure minimized the difference between the *χ*^2^ of the distribution of RTs in each cue-coherence-ISI bin and the RT distribution generated by the chosen model at the given parameters. Fitting was performed using a genetic algorithm (MATLAB Global Optimization Toolbox function ga) that ran for 1,000 generations per parameter, at a population size of 50 per parameter.

### Imaging analysis

To identify neural markers of stimulus reinstatement, we first defined patterns of activity in ventral visual stream regions that indicated participants were processing “face” or “scene” photographs. We then analyzed the degree to which these patterns were present during the post-cue, pre-stimulus ISIs in Phase 4. Because no pictures were present on-screen during this period of interest, we reasoned that greater evidence of stimulus reinstatement would indicate that participants were recalling the cued photograph. We therefore predicted that this reinstatement evidence would be reflected in response accuracies, response times, and DDM model parameters.

#### ROI definition

We identified a region of interest consisting of voxels that (across the group) showed preferential activation to face or scene photographs, using the following procedure.

First, for each participant, we performed a GLM analysis of BOLD signal during the localizer task. We identified voxels that responded more to scenes or faces, relative to other categories (univariate contrasts: faces *>* scenes | scrambled scenes | objects; scenes *>* faces | scrambled scenes | objects). For each participant, we selected clusters in the posterior parahippocampal region (matching the reported Parahippocampal Place Area (PPA); [26]) and posterior fusiform gyrus (matching the reported Fusiform Face Area (FFA); [25]) that were significant at p < 0.005, uncorrected. Next, each per-participant voxel mask was binarized; all above-threshold voxels were set to 1. To regularize the ROIs and ensure they were consistent across participants, the resulting individual mask was then warped to match the group average anatomical; these group-space masks were added together and the summed image thresholded to include all voxels present in more than 90% of participants. This final group ROI was then warped back to the individual participant space, and the result used as the final mask for pattern analysis.

#### Stimulus-specific pattern analysis

We computed the pattern of activity for each target photograph, across the corresponding category-preferring ROI. For each photograph in each block, we took the average pattern of activity over the last five presentations of the photograph during Phase 1. (The first five presentations were excluded to allow repetition suppression and learning effects to stabilize.)

We next used these four patterns as a template for analyzing activity during the post-cue, pre-stimulus ISI in Phase 4. For each trial, we computed, within the ROI corresponding to the cued category, the pattern of activity between the time of cue onset and either the time of response or one TR before the onset of the flickering stream, whichever came first. We then correlated this activity pattern with the corresponding target pattern, defined above. These correlation values, one for each Phase 4 trial, were then Fisher-transformed and used as predictor variables in our analyses of interest. We refer to these values as the *Reinstatement index*.

## Notes

https://openneuro.org/datasets/ds001614/

http://aaron.bornstein.org/cv/pubs/2022_baftnc_biorxiv_supp.pdf

### Competing Interest Statement

The authors have declared no competing interest.

### Summary of Updates

Revisions as requested by peer reviewers - in particular, added Discussion and a re-framing away from an evidence accumulation-specific account. Also, individual datapoints were added to the relevant figures.

## References

[1] Antonio Rangel, Colin Camerer, and P Read Montague. A framework for studying the neurobiology of value-based decision making. Nature Reviews Neuroscience, 9(7):545–56, jul 2008. ISSN 1471-0048. doi: 10.1038/nrn2357. URL http://www.ncbi.nlm.nih.gov/pubmed/18545266.

[2] Joshua I Gold and Michael N Shadlen. The neural basis of decision making. Annual Review of Neuroscience, 30:535–74, jan 2007. ISSN 0147-006X. doi: 10.1146/annurev.neuro.29.051605.113038. URL http://www.ncbi.nlm.nih.gov/pubmed/17600525.

[3] Roger Ratcliff. A Theory of Memory Retrieval. Psychological Review, 85(2):59–108, 1978.

[4] Alan M Gordon, Jesse Rissman, Roozbeh Kiani, and Anthony D Wagner. Cortical Reinstatement Mediates the Relationship Between Content-Specific Encoding Activity and Subsequent Recollection Decisions. Cerebral Cortex, 24(12): 3350–3364, aug 2013. ISSN 1460-2199. doi: 10.1093/cercor/bht194. URL http://www.ncbi.nlm.nih.gov/pubmed/23921785.

[5] Don van Ravenzwaaij, Martijn J Mulder, Francis Tuerlinckx, and Eric-Jan Wagenmakers. Do the dynamics of prior information depend on task context? An analysis of optimal performance and an empirical test. Frontiers in psychology, 3(May):132, jan 2012. ISSN 1664-1078. doi: 10.3389/fpsyg.2012.00132.

[6] Jan Drugowitsch, Rubén Moreno-Bote, Anne K Churchland, Michael N Shadlen, and Alexandre Pouget. The cost of accumulating evidence in perceptual decision making. Journal of Neuroscience, 32(11):3612–28, mar 2012. ISSN 1529-2401. doi: 10.1523/JNEUROSCI.4010-11.2012. URL http://www.ncbi.nlm.nih.gov/pubmed/22423085.

[7] Rani Moran. Optimal decision making in heterogeneous and biased environments. Psychonomic Bulletin & Review, pages 38–53, 2015. doi: 10.3758/s13423-014-0669-3.

[8] Don Van Ravenzwaaij, Martijn J Mulder, Francis Tuerlinckx, and Eric Jan Wagenmakers. Paradoxes of Optimal Decision Making : A Response to Moran (2015). Psychonomic Bulletin & Review, 22:307–308, 2015.

[9] Alessandra Perugini, Jochen Ditterich, and Michele A. Basso. Patients with Parkinson’s Disease Show Impaired Use of Priors in Conditions of Sensory Uncertainty. Current Biology, 26(14):1902–1910, 2016. ISSN 09609822. doi: 10.1016/j.cub.2016.05.039.

[10] Alessandra Perugini and Michele A Basso. Perceptual Decisions Based on Previously Learned Information are Independent of Dopaminergic Tone. Journal of Neurophysiology, page jn.00761.2017, 2017. ISSN 0022-3077. doi: 10.1152/jn.00761.2017. URL http://www.physiology.org/doi/10.1152/jn.00761.2017.

[11] Timothy D. Hanks, Mark E. Mazurek, Roozbeh Kiani, Elizabeth Hopp, N Michael, and M. N. Shadlen. Elapsed Decision Time Affects the Weighting of Prior Probability in a Perceptual Decision Task. Journal of Neuroscience, 31(17):6339–6352, apr 2011. ISSN 0270-6474. doi: 10.1523/JNEUROSCI.5613-10.2011. URL http://www.jneurosci.org/cgi/doi/10.1523/JNEUROSCI.5613-10.2011.

[12] Natalie Biderman, Akram Bakkour, and Daphna Shohamy. What are memories for? the hippocampus bridges past experience with future decisions. Trends in Cognitive Sciences, 24(7):542–556, 2020.

[13] Aaron M. Bornstein and Nathaniel D. Daw. Cortical and Hippocampal Correlates of Deliberation During Model-Based Decisions for Rewards in Humans. PLoS Computational Biology, 9(12):e1003387, dec 2013. ISSN 1553-7358. doi: 10.1371/journal.pcbi.1003387. URL http://dx.plos.org/10.1371/journal.pcbi.1003387.

[14] Michael N Shadlen and Daphna Shohamy. Decision Making and Sequential Sampling from Memory. Neuron, 90(5):927–939, 2016. ISSN 0896-6273. doi: 10.1016/j.neuron.2016.04.036. URL http://dx.doi.org/10.1016/j.neuron.2016.04.036.

[15] Aaron M Bornstein, Mel W Khaw, Daphna Shohamy, and Nathaniel D. Daw. Reminders of past choices bias decisions for reward in humans. Nature Communications, 8:15958, 2017. doi: 10.1038/ncomms15958.

[16] Aaron M Bornstein and Kenneth A Norman. Reinstated episodic context guides sampling-based decisions for reward. Nature Neuroscience, 2017. doi: 10.1038/nn.4573.

[17] Akram Bakkour, Daniela J Palombo, Ariel Zylberberg, Yul HR Kang, Allison Reid, Mieke Verfaellie, Michael N Shadlen, and Daphna Shohamy. The hippocampus supports deliberation during value-based decisions. elife, 8:e46080, 2019.

[18] Aaron M. Bornstein and Nathaniel D. Daw. Dissociating hippocampal and striatal contributions to sequential prediction learning. European Journal of Neuroscience, 35 (7):1011–1023, apr 2012. ISSN 0953816X. doi: 10.1111/j.1460-9568.2011.07920.x. URL http://doi.wiley.com/10.1111/j.1460-9568.2011.07920.x.

[19] Kim S Graham, Morgan D Barense, and Andy CH Lee. Going beyond ltm in the mtl: a synthesis of neuropsychological and neuroimaging findings on the role of the medial temporal lobe in memory and perception. Neuropsychologia, 48(4):831–853, 2010.

[20] Nicolette Sullivan, Cendri Hutcherson, Alison Harris, and Antonio Rangel. Dietary self-control is related to the speed with which attributes of healthfulness and tastiness are processed. Psychological science, 26(2):122–134, 2015.

[21] Vaibhav Srivastava, Samuel F. Feng, Jonathan D. Cohen, Naomi Ehrich Leonard, and Amitai Shenhav. A martingale analysis of first passage times of time-dependent Wiener diffusion models. Journal of Mathematical Psychology, 2016. ISSN 00222496. doi: 10.1016/j.jmp.2016.10.001. URL http://dx.doi.org/10.1016/j.jmp.2016.10.001.

[22] Gaia Lombardi and Todd Hare. Piecewise constant averaging methods allow for fast and accurate hierarchical bayesian estimation of drift diffusion models with time-varying evidence accumulation rates. 2021.

[23] J. a. Hartigan and P. M. Hartigan. The Dip Test of Unimodality. The Annals of Statistics, 13(1):70–84, 1985. ISSN 0090-5364. doi: 10.1214/aos/1176346577.

[24] Rafal Bogacz, Eric Brown, Jeff Moehlis, Philip Holmes, and Jonathan D Cohen. The physics of optimal decision making: a formal analysis of models of performance in two-alternative forced-choice tasks. Psychological Review, 113(4): 700–65, oct 2006. ISSN 0033-295X. doi: 10.1037/0033-295X.113.4.700. URL http://www.ncbi.nlm.nih.gov/pubmed/17014301.

[25] Nancy Kanwisher, Josh Mcdermott, and Marvin M Chun. The Fusiform Face Area: A Module in Human Extrastriate Cortex Specialized for Face Perception. Journal of Neuroscience, 17(11):4302–4311, 1997.

[26] R Epstein and N Kanwisher. A cortical representation of the local visual environment. Nature, 392(6676):598–601, apr 1998. ISSN 0028-0836. doi: 10.1038/33402. URL http://www.ncbi.nlm.nih.gov/pubmed/9560155.

[27] Rafal Bogacz, Peter T Hu, Philip J Holmes, and Jonathan D Cohen. Do humans produce the speed-accuracy trade-off that maximizes reward rate? Quarterly Journal of Experimental Psychology, 63(5):863–91, 2010. ISSN 1747-0226. doi: 10.1080/17470210903091643.

[28] C. M. Bishop. Pattern Recognition and Machine Learning. Springer, London, 2006.

[29] Kyle E. Dunovan, Joshua J. Tremel, and Mark E. Wheeler. Prior probability and feature predictability interactively bias perceptual decisions. Neuropsychologia, 61(1): 210–221, jun 2014. ISSN 18733514. doi: 10.1016/j.neuropsychologia.2014.06.024. URL http://linkinghub.elsevier.com/retrieve/pii/S002839321400205X http://www.ncbi.nlm.nih.gov/pubmed/24978303.

[30] James J Palestro, Emily Weichart, B Sederberg, and Brandon M Turner. Some task demands induce collapsing bounds: Evidence from a behavioral analysis. Psychonomic Bulletin & Review, 2018. doi: 10.3758/s13423-018-1479-9.

[31] Fábio P Leite and Roger Ratcliff. What cognitive processes drive response biases ? A diffusion model analysis. Judgement and Decision Making, 6(7):651–687, 2011.

[32] Roger Ratcliff and Francis Tuerlinckx. Estimating parameters of the diffusion model: approaches to dealing with contaminant reaction times and parameter variability. Psychonomic bulletin & review, 9(3):438–81, sep 2002. ISSN 1069-9384. URL http://www.pubmedcentral.nih.gov/articlerender.fcgi?artid=2474747&tool=pmcentrez&ren

[33] Roger Ratcliff and Gail McKoon. The diffusion decision model: theory and data for two-choice decision tasks. Neural computation, 20(4):873–922, apr 2008. ISSN 0899-7667. doi: 10.1162/neco.2008.12-06-420. URL http://www.pubmedcentral.nih.gov/articlerender.fcgi?artid=2474742&tool=pmcentrez&ren

[34] Christopher K Hauser, Dantong Zhu, Terrence R Stanford, and Emilio Salinas. Motor selection dynamics in FEF explain the reaction time variance of saccades to single targets. eLife, pages 1–32, 2018.

[35] Sophie Deneve. Making decisions with unknown sensory reliability. Frontiers in neuroscience, 6:75, 2012.

[36] D. Rahnev, H. Lau, and F. P. de Lange. Prior Expectation Modulates the Interaction between Sensory and Prefrontal Regions in the Human Brain. Journal of Neuroscience, 31 (29):10741–10748, jul 2011. ISSN 0270-6474. doi: 10.1523/JNEUROSCI.1478-11.2011. URL http://www.jneurosci.org/cgi/doi/10.1523/JNEUROSCI.1478-11.2011.

[37] Valentin Wyart, Vincent deGardelle, Jacqueline Scholl, Christopher Summerfield, Vincent De Gardelle, Jacqueline Scholl, and Christopher Summerfield. Rhythmic Fluctuations in Evidence Accumulation during Decision Making in the Human Brain. Neuron, 76(4):847–858, nov 2012. ISSN 08966273. doi: 10.1016/j.neuron.2012.09.015. URL http://linkinghub.elsevier.com/retrieve/pii/S0896627312008483.

[38] Angela J Yu and Jonathan D Cohen. Sequential effects: Superstition or rational behavior? Advances in Neural Information Processing Systems, 21:1873–80, 2009. ISSN 1049-5258 (Print). URL http://www.pni.princeton.edu/ncc/publications/2009/YuNIPS09.pdf.

[39] David Silver and Joel Veness. Monte-Carlo Planning in Large POMDPs. Neural Information Processing Systems, pages 1–9, 2010.

[40] Andrew C Barto and Richard S Sutton. Reinforcement Learning: An Introduction. MIT Press, Cambridge, MA, apr 1998. doi: 10.1007/s00134-010-1760-5.

[41] Mate Lengyel and Peter Dayan. Hippocampal Contributions to Control: The Third Way. Advances in Neural Information Processing Systems, 20:889–896, 2008.

[42] Samuel Ritter, Jane X Wang, Zeb Kurth-nelson Siddhant, Charles Blundell, Razvan Pascanu, and Matthew Botvinick. Been There, Done That: Meta-Learning with Episodic Recall. In ICML, 2018.

[43] Oliver Vikbladh, Daphna Shohamy, and Nathaniel D Daw. Episodic Contributions to Model - Based Reinforcement Learning. Cognitive Computational Neuroscience Conference, pages 2016–2017, 2017.

[44] Roozbeh Kiani, Anne K Churchland, and Michael N Shadlen. Integration of Direction Cues Is Invariant to the Temporal Gap between Them. Journal of Neuroscience, 33 (42):16483–16489, 2013. doi: 10.1523/JNEUROSCI.2094-13.2013.

[45] Roozbeh Kiani, Timothy D Hanks, and Michael N Shadlen. Bounded Integration in Parietal Cortex Underlies Decisions Even When Viewing Duration Is Dictated by the Environment. Journal of Neuroscience, 28(12):3017–3029, 2008. doi: 10.1523/JNEUROSCI.4761-07.2008.

[46] Abigail N Hoskin, Aaron M Bornstein, Kenneth A Norman, and Jonathan D Cohen. Refresh my memory: Episodic memory reinstatements intrude on working memory maintenance. Cognitive, Affective, & Behavioral Neuroscience, 19(2):338–354, 2019.

[47] Juan Gao, Rebecca Tortell, and James L McClelland. Dynamic integration of reward and stimulus information in perceptual decision-making. PloS one, 6(3): e16749, jan 2011. ISSN 1932-6203. doi: 10.1371/journal.pone.0016749. URL http://www.pubmedcentral.nih.gov/articlerender.fcgi?artid=3048391&tool=pmcentrez&ren

[48] X Anke Braun, X Anne E Urai, and X Tobias H Donner. Adaptive History Biases Result from Confidence-Weighted Accumulation of past Choices. Journal of Neuroscience, 38 (10):2418–2429, 2018. doi: 10.1523/JNEUROSCI.2189-17.2017.

[49] Brian Odegaard, Piercesare Grimaldi, Seong Hah, Megan A K Peters, and Hakwan Lau. Superior colliculus neuronal ensemble activity signals optimal rather than subjective confidence. Proceedings of the National Academy of Sciences, pages 1–10, 2017. doi: 10.1073/pnas.1711628115.

[50] Roozbeh Kiani and Michael N Shadlen. Representation of confidence associated with a decision by neurons in the parietal cortex. Science, 324(5928): 759–64, may 2009. ISSN 1095-9203. doi: 10.1126/science.1169405. URL http://www.ncbi.nlm.nih.gov/pubmed/19423820.

[51] Arbora Resulaj, Roozbeh Kiani, Daniel M Wolpert, and Michael N Shadlen. Changes of mind in decision-making. Nature, 461(7261):263–6, 2009. ISSN 1476-4687. doi: 10.1038/nature08275. URL http://www.ncbi.nlm.nih.gov/pubmed/19693010.

[52] Ronald vandenBerg, Ariel Zylberberg, Roozbeh Kiani, MichaelN. Shadlen, DanielM. Wolpert, S.W. Link, R. Ratcliff, J.N. Rouder, D. Thura, J. Beauregard-Racine, C.W. Fradet, P. Cisek, J. Drugowitsch, R. Moreno-Bote, A.K. Churchland, M.N. Shadlen, A. Pouget, J. Drugowitsch, G.C. DeAngelis, D.E. Angelaki, A. Pouget, B.A. Purcell, R. Kiani, D. Laming, R.P. Heitz, J.D. Schall, N. Kolling, T.E. Behrens, R.B. Mars, M.F. Rushworth, B.B. Averbeck, R. Kiani, M.N. Shadlen, R. Kiani, L. Corthell, M.N. Shadlen, R. van den Berg, K. Anandalingam, A. Zylberberg, R. Kiani, M.N. Shadlen, D.M. Wolpert, B.W. Brunton, M.M. Botvinick, C.D. Brody, J.I. Gold, M.N. Shadlen, M.N. Shadlen, R. Kiani, A. Resulaj, R. Kiani, D.M. Wolpert, M.N. Shadlen, D. Burk, J.N. Ingram, D.W. Franklin, M.N. Shadlen, D.M. Wolpert, J. Moher, J.H. Song, C.R. Fetsch, R. Kiani, W.T. Newsome, M.N. Shadlen, S. Yu, T.J. Pleskac, M.D. Zeigenfuse, T.J. Pleskac, J.R. Busemeyer, J. Palmer, A.C. Huk, M.N. Shadlen, P.L. Smith, R. Ratcliff, A. Pouget, J. Drugowitsch, A. Kepecs, R.E. Kass, A.E. Raftery, H. Jeffreys, Y. Huang, T. Hanks, M. Shadlen, A.L. Friesen, R.P. Rao, D. Laming, M.R. Cohen, W.T. Newsome, J.F. Mitchell, K.A. Sundberg, J.H. Reynolds, M.R. Cohen, J.H. Maunsell, J.D. Roitman, M.N. Shadlen, T. Hanks, R. Kiani, M.N. Shadlen, A.K. Churchland, R. Kiani, M.N. Shadlen, M.E. Mazurek, J.D. Roitman, J. Ditterich, M.N. Shadlen, M. Usher, J.L. McClelland, C.C. Lo, X.J. Wang, L. Ding, J.I. Gold, L. Ding, J.I. Gold, A. Kepecs, N. Uchida, H.A. Zariwala, Z.F. Mainen, R.R. Hampton, D.H. Brainard, F.W. Cornelissen, E.M. Peters, J. Palmer, M.N. Shadlen, W.T. Newsome, and R. Ratcliff. Confidence Is the Bridge between Multi-stage Decisions. Current Biology, 0(0):114–135, 2016. ISSN 09609822. doi: 10.1016/j.cub.2016.10.021. URL http://linkinghub.elsevier.com/retrieve/pii/S0960982216312064.

[53] Li Yan Mccurdy, Brian Maniscalco, Janet Metcalfe, Ka Yuet Liu, Floris P De Lange, and Hakwan Lau. Anatomical Coupling between Distinct Metacognitive Systems for Memory and Visual Perception. Journal of Neuroscience, 33(5):1897–1906, 2013. doi: 10.1523/JNEUROSCI.1890-12.2013.

[54] Qun Ye, Futing Zou, Hakwan Lau, Yi Hu, and Sze Chai Kwok. Causal Evidence for Mnemonic Metacognition in Human Precuneus. Journal of Neuroscience, 38(28):6379–6387, 2018. doi: 10.1523/JNEUROSCI.0660-18.2018.

[55] Ari Khoudary, Megan AK Peters, and Aaron M Bornstein. Precision-weighted evidence integration predicts time-varying influence of memory on perceptual decisions. Cognitive Computational Neuroscience, 2022.

[56] Shaoming Wang, Samuel F Feng, and Aaron M Bornstein. Mixing memory and desire: How memory reactivation supports deliberative decision-making. Wiley Interdisciplinary Reviews: Cognitive Science, 13(2):e1581, 2022.

[57] Xiaochen Y Zheng, Martin N Hebart, Raymond J Dolan, Christian F Doeller, Roshan Cools, and Mona M Garvert. Parallel cognitive maps for short-term statistical and long-term semantic relationships in the hippocampal formation. bioRxiv, pages 2022–08, 2022.

[58] Gabriel M Stine, Ariel Zylberberg, Jochen Ditterich, and Michael N Shadlen. Differentiating between integration and non-integration strategies in perceptual decision making. Elife, 9:e55365, 2020.

[59] Claire M. Gillan, Michal Kosinski, Robert Whelan, Elizabeth A. Phelps, and Nathaniel D. Daw. Characterizing a psychiatric symptom dimension related to deficits in goaldirected control. eLife, 5(MARCH2016):1–24, 2016. ISSN 2050084X. doi: 10.7554/eLife.11305.

[60] Daniel J Davies, Christoph Teufel, and Paul C Fletcher. Anomalous Perceptions and Beliefs Are Associated With Shifts Toward Different Types of Prior Knowledge in Perceptual Inference. Schizophrenia Bulletin, (January), 2017. ISSN 0586-7614. doi: 10.1093/schbul/sbx177. URL http://academic.oup.com/schizophreniabulletin/advance-article/doi/10.1093/schbul/sbx

[61] D H Brainard. The Psychophysics Toolbox. Spatial Vision, 10(4):433–6, jan 1997. ISSN 0169-1015. URL http://www.ncbi.nlm.nih.gov/pubmed/9176952.

[62] A. B. Watson and Dennis G. Pelli. QUEST: A Bayesian adaptive psychometric method. Perception & Psychophysics, 33(2):113–120, 1983.

[63] Stephen M. Smith, Mark Jenkinson, Mark W. Woolrich, Christian F. Beckmann, Timothy E J Behrens, Heidi Johansen-Berg, Peter R. Bannister, Marilena De Luca, Ivana Drobnjak, David E. Flitney, Rami K. Niazy, James Saunders, John Vickers, Yongyue Zhang, Nicola De Stefano, J. Michael Brady, and Paul M. Matthews. Advances in functional and structural MR image analysis and implementation as FSL. NeuroImage, 23 (SUPPL. 1):208–219, 2004. ISSN 10538119. doi: 10.1016/j.neuroimage.2004.07.051.

[64] Mark Jenkinson, Christian F. Beckmann, Timothy E J Behrens, Mark W. Woolrich, and Stephen M. Smith. FSL. NeuroImage, 62(2):782–790, 2012. ISSN 10538119. doi: 10.1016/j.neuroimage.2011.09.015.

[65] Nicholas Price and F. Mechler. HartigansDipTest, 2002. URL http://www.nicprice.net/diptest/.

[66] Ghootae Kim, Jarrod A Lewis-peacock, Kenneth A Norman, and Nicholas B Turk-browne. Pruning of memories by context-based prediction error. Proceedings of the National Academy of Sciences, 111(24), 2014. doi: 10.1073/pnas.1319438111.

[67] Jacob Cohen. Statistical power analysis for the behavioural sciences. NJ: Lawrence Earlbaum Associates, Hillside, NJ, 1988.

