## Supplemental material for "Associative memory retrieval modulates upcoming perceptual decisions"

### Model fits

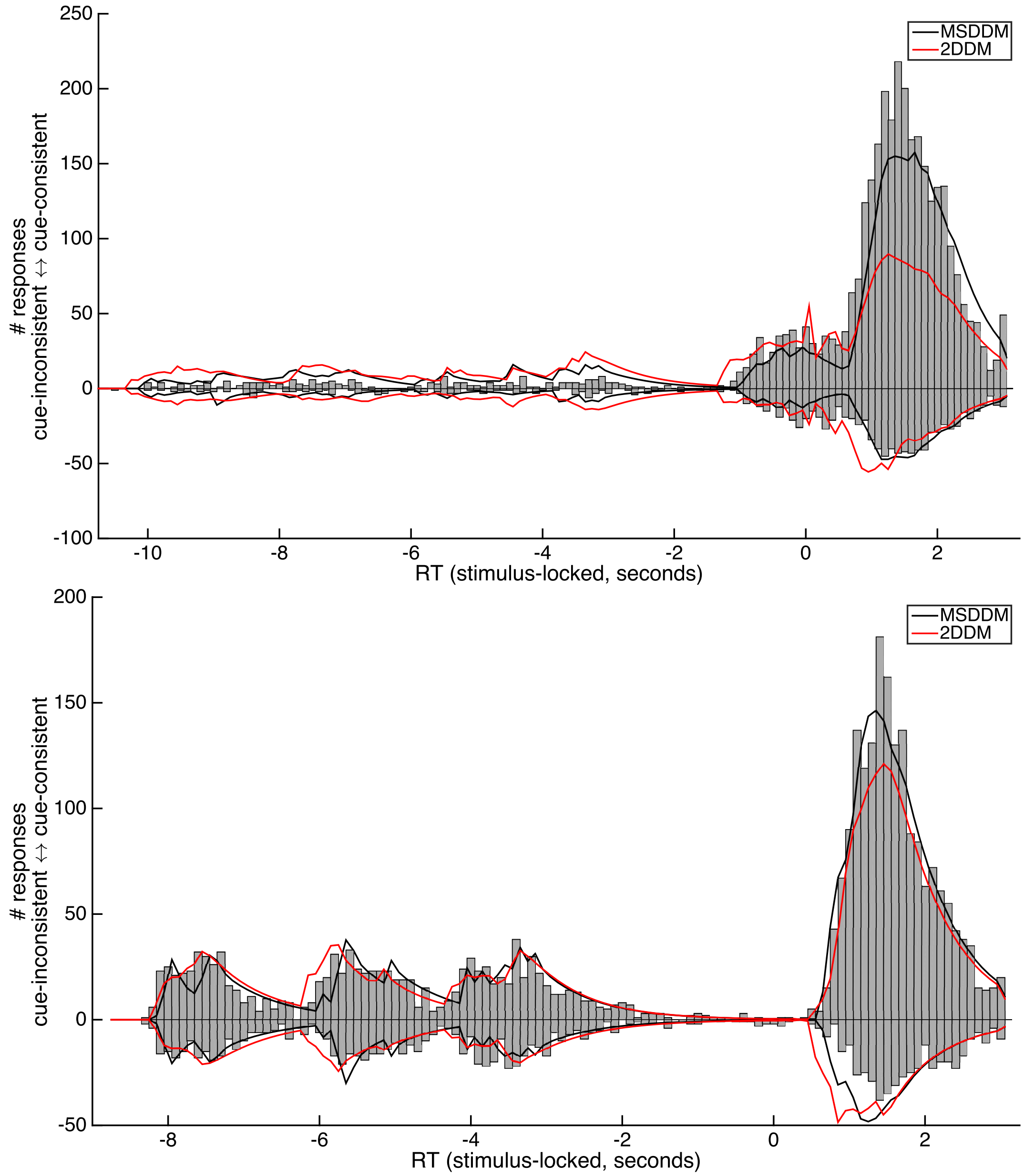


Figure S1: **MSDDM & 2DDM model fits overlaid on RT histograms**.

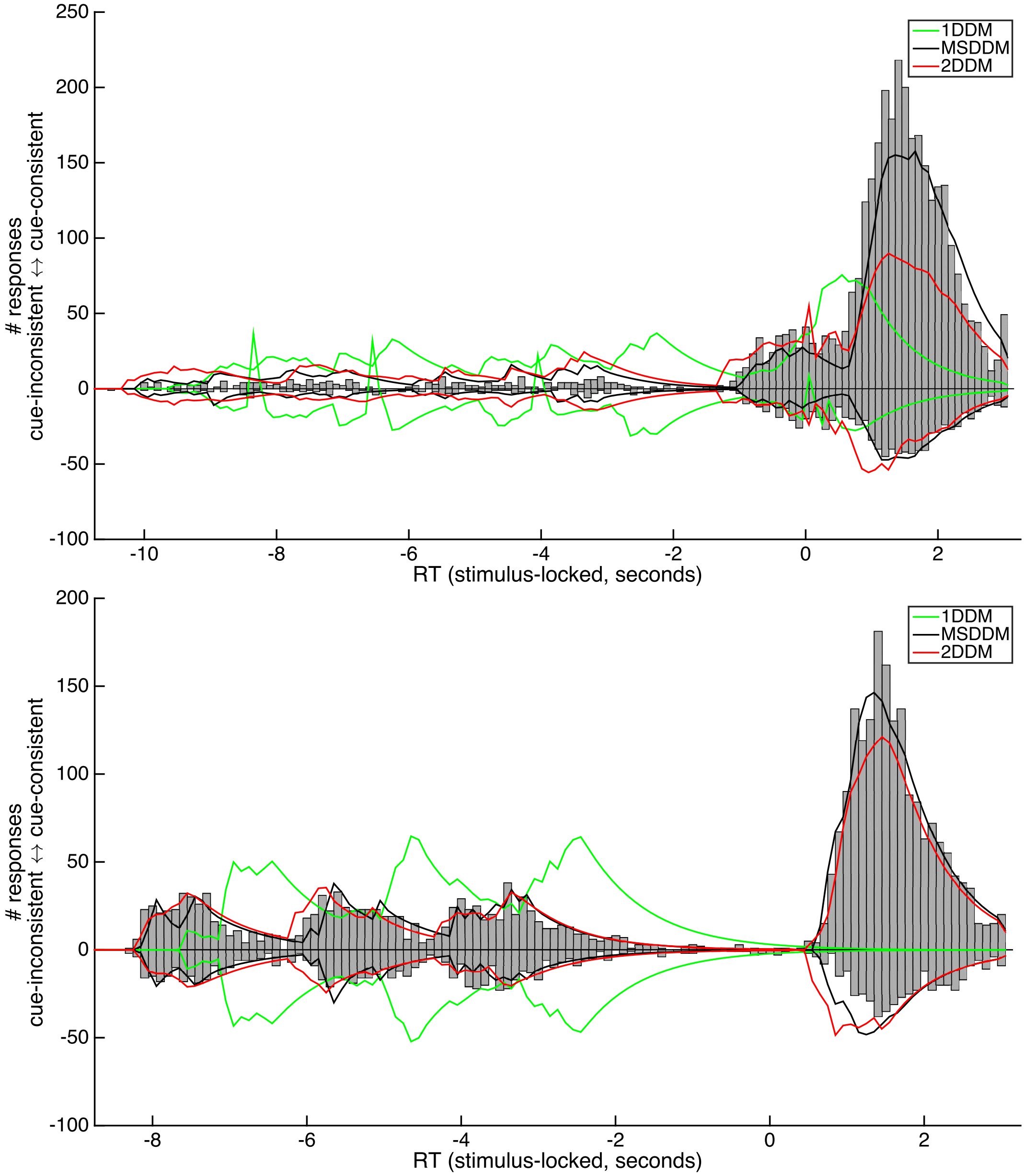


Figure S2: **Model fits overlaid on RT histograms, with addition of 1DDM.**

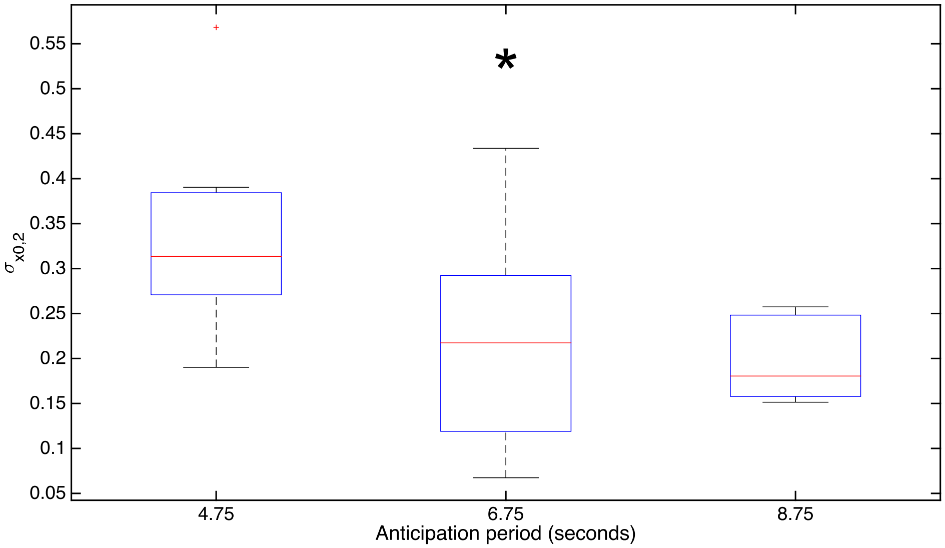


Figure S3: **Second-stage starting point variability decreases with ISI**.

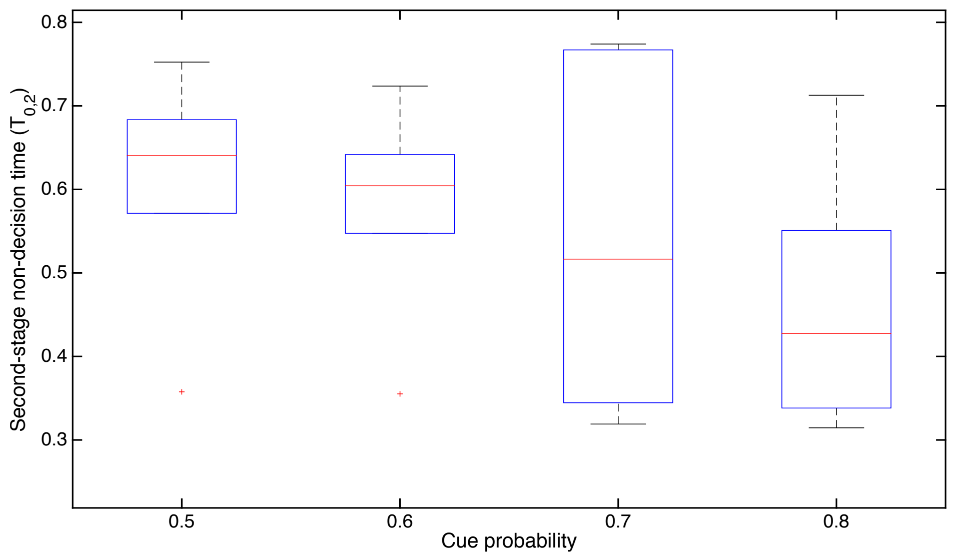


Figure S4: **Second-stage non-decision time decreases with cue probability**.

| cue | coh | ISI | $\eta_1$ | $\eta_2$ | $x_{0,1}$ | $\sigma_{x_{0,1}}$ | $x_{0,2}$ | $\sigma_{x_{0,2}}$ | $T_{0,1}$ | $T_{0,2}$ |
|---|---|---|---|---|---|---|---|---|---|---|
| 0.50 | 0.65 | 0.50 | -0.06 | 0.49 | 0.00 | 0.22 | -0.08 | 0.33 | 0.59 | 0.71 |
| | | 1.00 | 2.03 | 0.39 | -0.06 | 1.36 | -0.18 | 0.28 | 0.44 | 0.48 |
| | | 4.00 | -0.08 | 0.49 | -0.14 | 0.34 | -0.19 | 0.29 | 0.82 | 0.79 |
| | | 6.00 | 0.05 | 0.43 | 0.12 | 0.69 | -0.35 | 0.09 | 0.46 | 0.31 |
| | | 8.00 | -0.06 | 0.37 | 0.25 | 0.50 | 0.13 | 0.25 | 0.93 | 0.47 |
| | | 10.00 | -0.05 | 0.18 | 0.04 | 0.37 | -0.35 | 1.30 | 0.42 | 0.27 |
| | 0.85 | 0.50 | 0.19 | 0.95 | -0.08 | 0.00 | -0.24 | 0.23 | 0.43 | 0.26 |
| | | 1.00 | 0.80 | 0.56 | -0.26 | 1.29 | -0.22 | 0.39 | 0.54 | 0.31 |
| | | 4.00 | 0.09 | 0.42 | -0.11 | 0.28 | 0.32 | 0.26 | 0.62 | 0.58 |
| | | 6.00 | 0.03 | 1.68 | -0.36 | 0.20 | -0.24 | 0.24 | 1.07 | 0.59 |
| | | 8.00 | 0.65 | 1.62 | -0.35 | 0.11 | -0.21 | 0.14 | 1.07 | 0.71 |
| | | 10.00 | -0.05 | 0.18 | 0.10 | 0.37 | -0.31 | 1.30 | 0.44 | 0.26 |
| 0.60 | 0.65 | 0.50 | 0.17 | 0.75 | 0.01 | 0.00 | -0.15 | 0.54 | 0.52 | 0.72 |
| | | 1.00 | 0.91 | 0.31 | -0.48 | 0.92 | -0.25 | 0.28 | 0.44 | 0.40 |
| | | 4.00 | 0.08 | 0.65 | 0.10 | 0.28 | -0.11 | 0.49 | 0.61 | 0.88 |
| | | 6.00 | 0.51 | 0.69 | 0.82 | 1.93 | -0.31 | 0.29 | 0.43 | 0.88 |
| | | 8.00 | 2.16 | 0.40 | -0.01 | 2.02 | -0.11 | 0.27 | 0.50 | 0.90 |
| | | 10.00 | 1.35 | 0.04 | 0.27 | 1.76 | 0.01 | 1.00 | 0.42 | 1.02 |
| | 0.85 | 0.50 | 0.34 | 0.88 | -0.13 | 0.38 | -0.20 | 0.15 | 0.77 | 0.33 |
| | | 1.00 | 0.69 | 0.48 | -0.07 | 1.51 | -0.22 | 0.55 | 0.58 | 0.29 |
| | | 4.00 | 0.26 | 0.37 | 0.07 | 0.66 | -0.37 | 1.15 | 0.47 | 0.35 |
| | | 6.00 | 0.21 | 0.94 | -0.08 | 0.03 | -0.00 | 0.45 | 1.10 | 0.46 |
| | | 8.00 | 0.06 | 0.35 | 0.42 | 1.67 | -0.64 | 1.17 | 0.98 | 0.42 |
| | | 10.00 | -0.06 | 0.18 | 0.05 | 0.35 | -0.36 | 1.30 | 0.42 | 0.26 |
| 0.70 | 0.65 | 0.50 | 1.05 | 0.24 | 0.03 | 1.35 | -0.17 | 0.83 | 0.63 | 0.30 |
| | | 1.00 | 0.95 | 0.36 | 0.10 | 0.94 | -0.20 | 0.38 | 0.46 | 0.32 |
| | | 4.00 | -0.09 | 0.70 | 0.24 | 0.34 | -0.07 | 0.43 | 0.90 | 0.58 |
| | | 6.00 | -0.08 | 0.74 | 0.09 | 0.24 | -0.03 | 0.55 | 0.90 | 0.65 |
| | | 8.00 | -0.10 | 0.10 | 0.09 | 0.25 | 0.14 | 0.28 | 1.03 | 0.57 |
| | | 10.00 | 0.07 | 0.21 | -0.01 | 0.37 | -0.36 | 1.43 | 0.43 | 0.27 |
| | 0.85 | 0.50 | 0.53 | 1.03 | -0.05 | 0.12 | -0.02 | 0.49 | 0.50 | 0.64 |
| | | 1.00 | 1.11 | 0.42 | -0.56 | 0.77 | -0.20 | 0.33 | 0.45 | 0.35 |
| | | 4.00 | 1.00 | 1.31 | 0.19 | 0.92 | -0.34 | 0.40 | 0.60 | 0.48 |
| 0.80 | 0.65 | 0.50 | 1.00 | 0.21 | 0.08 | 0.72 | -0.09 | 0.98 | 0.49 | 0.30 |
| | | 1.00 | 1.26 | 0.30 | 0.07 | 0.53 | -0.26 | 1.24 | 0.44 | 0.38 |
| | | 4.00 | 0.27 | 0.72 | 0.22 | 0.54 | -0.35 | 0.93 | 0.42 | 0.31 |
| | | 6.00 | 1.65 | 1.32 | 0.24 | 1.11 | -0.19 | 1.23 | 1.01 | 0.90 |
| | | 8.00 | 0.71 | 0.20 | -0.05 | 0.26 | -0.33 | 0.19 | 0.71 | 0.81 |
| | | 10.00 | -0.06 | 0.18 | 0.08 | 0.35 | -0.36 | 1.30 | 0.42 | 0.26 |
| | 0.85 | 0.50 | 1.24 | 0.41 | -0.53 | 0.54 | -0.03 | 0.62 | 0.46 | 0.33 |
| | | 1.00 | 1.01 | 0.91 | -0.23 | 0.84 | -0.30 | 0.34 | 0.54 | 0.35 |
| | | 4.00 | 0.30 | 0.21 | 0.19 | 0.68 | 0.24 | 0.49 | 0.78 | 0.23 |
| | | 6.00 | -0.09 | 0.98 | 0.64 | 0.55 | -0.08 | 0.31 | 0.92 | 0.58 |
| | | 8.00 | 0.64 | 0.76 | 0.01 | 0.29 | -0.09 | 0.48 | 0.57 | 0.47 |
| | | 10.00 | 0.69 | 0.35 | 0.14 | 1.04 | -0.32 | 1.05 | 0.42 | 0.78 |

Table S1: **Experiment 1: Parameter fits for two, unconnected DDMs (2DDM)**.

| cue | coh | ISI | $\eta_1$ | $\eta_2$ | $x_{0,1}$ | $\sigma_{x_{0,1}}$ | $T_{0,1}$ |
|---|---|---|---|---|---|---|---|
| 0.50 | 0.65 | 0.50 | 0.26 | 0.47 | 0.96 | 4.99 | 1.02 |
| | | 1.00 | -0.06 | 0.47 | 0.98 | 4.95 | 0.97 |
| | | 4.00 | -0.07 | 0.44 | -0.88 | 3.94 | 1.05 |
| | | 6.00 | 0.71 | 0.35 | 0.39 | 2.09 | 1.01 |
| | | 8.00 | 0.36 | 0.39 | 0.37 | 3.03 | 0.58 |
| | | 10.00 | 0.08 | 0.50 | 0.19 | 3.20 | 0.55 |
| | 0.85 | 0.50 | 0.00 | 0.94 | 0.19 | 3.20 | 0.49 |
| | | 1.00 | 0.47 | 0.79 | 0.17 | 2.85 | 0.59 |
| | | 4.00 | -0.06 | 0.88 | 0.97 | 3.00 | 0.50 |
| | | 6.00 | 0.31 | 2.16 | 0.99 | 4.98 | 0.89 |
| | | 8.00 | 1.40 | 1.24 | 0.15 | 2.95 | 0.91 |
| | | 10.00 | 0.09 | 1.20 | 0.19 | 3.01 | 0.54 |
| 0.60 | 0.65 | 0.50 | 0.52 | 0.57 | 0.24 | 2.27 | 0.88 |
| | | 1.00 | 0.78 | 0.38 | 0.23 | 2.31 | 0.90 |
| | | 4.00 | 0.76 | 0.58 | 0.39 | 2.47 | 0.86 |
| | | 6.00 | -0.10 | 1.03 | -0.99 | 5.00 | 1.09 |
| | | 8.00 | -0.04 | 0.33 | -0.95 | 4.98 | 1.03 |
| | | 10.00 | 0.89 | 0.20 | 0.27 | 2.49 | 1.04 |
| | 0.85 | 0.50 | 1.23 | 0.84 | 0.03 | 1.98 | 0.89 |
| | | 1.00 | 0.57 | 0.91 | 0.08 | 1.97 | 0.71 |
| | | 4.00 | 0.12 | 0.56 | 0.98 | 4.85 | 0.60 |
| | | 6.00 | 0.22 | 0.89 | 0.23 | 3.97 | 0.45 |
| | | 8.00 | 0.15 | 1.10 | 0.15 | 3.36 | 0.56 |
| | | 10.00 | 0.02 | 1.14 | 0.17 | 3.18 | 0.46 |
| 0.70 | 0.65 | 0.50 | 0.75 | 0.59 | 0.17 | 2.42 | 0.80 |
| | | 1.00 | 0.59 | 0.54 | 0.17 | 2.03 | 0.72 |
| | | 4.00 | 0.17 | 0.50 | 0.18 | 2.13 | 0.51 |
| | | 6.00 | 0.30 | 0.71 | 0.18 | 2.36 | 0.69 |
| | | 8.00 | 0.09 | 0.25 | 0.94 | 4.93 | 0.71 |
| | | 10.00 | 0.27 | 0.46 | 0.17 | 3.20 | 0.55 |
| | 0.85 | 0.50 | 0.95 | 0.93 | -0.06 | 2.05 | 0.81 |
| | | 1.00 | 0.93 | 0.77 | 0.41 | 2.03 | 0.78 |
| | | 4.00 | 1.13 | 1.12 | 0.63 | 2.50 | 0.81 |
| 0.80 | 0.65 | 0.50 | 0.94 | 0.37 | 0.17 | 1.44 | 0.74 |
| | | 1.00 | 1.64 | 0.24 | -0.24 | 1.83 | 0.82 |
| | | 4.00 | 0.36 | 0.92 | 0.89 | 1.96 | 0.62 |
| | | 6.00 | 1.03 | 0.88 | 0.36 | 0.61 | 0.99 |
| | | 8.00 | 0.79 | -0.01 | 0.15 | 0.79 | 0.85 |
| | | 10.00 | 0.62 | 0.79 | -0.27 | 0.68 | 0.44 |
| | 0.85 | 0.50 | 0.54 | 1.24 | 0.99 | 3.46 | 0.56 |
| | | 1.00 | 1.12 | 0.69 | 0.23 | 2.41 | 0.82 |
| | | 4.00 | 0.78 | 0.78 | -1.00 | 1.97 | 0.65 |
| | | 6.00 | 0.64 | 0.92 | 0.80 | 2.13 | 0.84 |
| | | 8.00 | 1.45 | 0.87 | 0.21 | 2.59 | 0.98 |
| | | 10.00 | 0.63 | 0.73 | -0.01 | 2.94 | 0.73 |

Table S2: **Experiment 1: Parameter fits for the Multi-Stage DDM (MS-DDM)**.

| cue | coh | ISI | $\eta_1$ | $\eta_2$ | $x_{0,1}$ | $\sigma_{x_{0,1}}$ | $x_{0,2}$ | $\sigma_{x_{0,2}}$ | $T_{0,1}$ | $T_{0,2}$ |
|---|---|---|---|---|---|---|---|---|---|---|
| 0.50 | 0.65 | 4.00 | -0.04 | 0.37 | -0.14 | 0.53 | 0.13 | 0.39 | 0.54 | 0.75 |
| 0.50 | 0.65 | 6.00 | 0.02 | 0.44 | -0.02 | 0.28 | -0.07 | 0.07 | 0.45 | 0.57 |
| 0.50 | 0.65 | 8.00 | 0.12 | 0.27 | -0.06 | 0.33 | 0.12 | 0.15 | 0.48 | 0.68 |
| 0.50 | 0.85 | 4.00 | 0.01 | 0.73 | 0.03 | 0.32 | 0.06 | 0.19 | 0.91 | 0.68 |
| 0.50 | 0.85 | 6.00 | 0.09 | 0.44 | 0.24 | 0.72 | -0.34 | 0.14 | 0.46 | 0.36 |
| 0.50 | 0.85 | 8.00 | 0.03 | 1.04 | 0.15 | 0.24 | -0.21 | 0.16 | 0.80 | 0.61 |
| 0.60 | 0.65 | 4.00 | -0.02 | 0.44 | 0.10 | 0.43 | -0.07 | 0.25 | 0.52 | 0.61 |
| 0.60 | 0.65 | 6.00 | 0.08 | 0.39 | 0.43 | 0.72 | -0.07 | 0.10 | 0.41 | 0.59 |
| 0.60 | 0.65 | 8.00 | -0.07 | 0.28 | 0.07 | 0.58 | 0.18 | 0.24 | 0.59 | 0.72 |
| 0.60 | 0.85 | 4.00 | 0.25 | 1.19 | 0.09 | 1.58 | -0.36 | 0.32 | 0.47 | 0.36 |
| 0.60 | 0.85 | 6.00 | 0.33 | 0.56 | 0.19 | 1.33 | -0.17 | 0.30 | 0.48 | 0.55 |
| 0.60 | 0.85 | 8.00 | 0.04 | 0.84 | 0.13 | 0.64 | -0.10 | 0.25 | 0.54 | 0.64 |
| 0.70 | 0.65 | 4.00 | 0.00 | 0.54 | 0.18 | 0.21 | 0.02 | 0.29 | 0.59 | 0.77 |
| 0.70 | 0.65 | 6.00 | 0.16 | 0.46 | 0.24 | 0.85 | -0.35 | 0.22 | 0.53 | 0.43 |
| 0.70 | 0.65 | 8.00 | -0.05 | 0.59 | 0.13 | 0.26 | -0.01 | 0.26 | 0.57 | 0.77 |
| 0.70 | 0.85 | 4.00 | 0.71 | 1.29 | -0.07 | 0.82 | -0.34 | 0.38 | 0.44 | 0.34 |
| 0.70 | 0.85 | 6.00 | 0.25 | 0.72 | 0.03 | 1.02 | -0.27 | 0.21 | 0.50 | 0.32 |
| 0.70 | 0.85 | 8.00 | 0.16 | 0.85 | 0.16 | 0.61 | 0.10 | 0.16 | 0.59 | 0.60 |
| 0.80 | 0.65 | 4.00 | 0.73 | 0.63 | 0.17 | 0.79 | -0.20 | 0.57 | 0.42 | 0.47 |
| 0.80 | 0.65 | 6.00 | 0.42 | 0.44 | 0.32 | 0.74 | -0.21 | 0.29 | 0.41 | 0.34 |
| 0.80 | 0.65 | 8.00 | 0.02 | 0.29 | 0.25 | 0.45 | 0.17 | 0.20 | 0.57 | 0.71 |
| 0.80 | 0.85 | 4.00 | 1.13 | 1.15 | -0.05 | 0.71 | -0.32 | 0.31 | 0.57 | 0.31 |
| 0.80 | 0.85 | 6.00 | 0.41 | 0.65 | 0.30 | 0.80 | -0.34 | 0.43 | 0.50 | 0.39 |
| 0.80 | 0.85 | 8.00 | 0.54 | 1.26 | 0.03 | 0.27 | -0.07 | 0.16 | 0.45 | 0.55 |

Table S3: **Experiment 2: Parameter fits for two, unconnected DDMs (2DDM)**.

| cue | coh | ISI | $\eta_1$ | $\eta_2$ | $x_{0,1}$ | $\sigma_{x_{0,1}}$ | $T_{0,1}$ |
|---|---|---|---|---|---|---|---|
| 0.50 | 0.65 | 4.00 | 0.09 | 0.46 | 0.20 | 1.80 | 0.59 |
| | | 6.00 | 0.12 | 0.27 | 0.39 | 1.77 | 0.65 |
| | | 8.00 | -0.00 | 0.40 | -0.38 | 1.64 | 0.60 |
| | 0.85 | 4.00 | 0.40 | 0.69 | 0.06 | 1.91 | 0.90 |
| | | 6.00 | 0.08 | 0.64 | 0.29 | 2.35 | 0.83 |
| | | 8.00 | -0.08 | 1.12 | 0.98 | 2.40 | 0.76 |
| 0.60 | 0.65 | 4.00 | 0.34 | 0.28 | 0.25 | 1.78 | 0.62 |
| | | 6.00 | 0.11 | 0.32 | 0.34 | 1.83 | 0.65 |
| | | 8.00 | -0.04 | 0.47 | 0.35 | 1.71 | 0.64 |
| | 0.85 | 4.00 | -0.08 | 1.06 | -0.98 | 2.51 | 0.64 |
| | | 6.00 | 0.66 | 0.82 | 0.16 | 2.45 | 0.77 |
| | | 8.00 | 0.25 | 0.76 | 0.10 | 3.42 | 0.59 |
| 0.70 | 0.65 | 4.00 | -0.04 | 0.60 | 0.90 | 2.08 | 0.91 |
| | | 6.00 | 0.45 | 0.49 | 0.44 | 2.12 | 0.88 |
| | | 8.00 | -0.06 | 0.74 | 0.96 | 1.71 | 0.99 |
| | 0.85 | 4.00 | 0.37 | 1.39 | 0.23 | 1.93 | 0.61 |
| | | 6.00 | 0.56 | 1.38 | -0.35 | 1.52 | 0.62 |
| | | 8.00 | 0.22 | 1.04 | 0.53 | 1.55 | 0.62 |
| 0.80 | 0.65 | 4.00 | 1.07 | 0.37 | 0.68 | 1.87 | 0.61 |
| | | 6.00 | 0.49 | 0.43 | 0.41 | 1.90 | 0.61 |
| | | 8.00 | 0.18 | 0.43 | 0.92 | 1.24 | 0.63 |
| | 0.85 | 4.00 | 0.77 | 1.26 | 0.42 | 0.99 | 0.62 |
| | | 6.00 | 0.63 | 0.84 | 0.10 | 1.10 | 0.61 |
| | | 8.00 | 0.92 | 0.94 | 0.32 | 1.34 | 0.62 |

Table S4: **Experiment 2: Parameter fits for the Multi-Stage DDM (MS-DDM)**.
